# Multiple Introductions of Highly Pathogenic Avian Influenza Viruses into the High Arctic: Svalbard and Jan Mayen, 2022–2025

**DOI:** 10.64898/2026.02.17.706283

**Authors:** Silje Granstad, Ragnhild Tønnessen, Bjørnar Ytrehus, Sebastien Descamps, Børge Moe, Hallvard Strøm, Geir Wing Gabrielsen, Sveinn Are Hanssen, Egil Rønning, Ole-Herman Tronerud, Maarten J.J.E. Loonen, Britt Gjerset, Hans Kristian Mjelde, Knut Madslien, Johan Åkerstedt, Cathrine Arnason Bøe

## Abstract

Between 2022 and 2025, highly pathogenic avian influenza viruses (HPAIVs) of clade 2.3.4.4b, including four distinct H5 Eurasian (EA) genotypes, were detected in wild birds and mammals in the Svalbard Archipelago and on the island of Jan Mayen. We describe their epidemiology and genomic characteristics to improve understanding of HPAIV occurrence and transmission in the High Arctic.

The initial cases in 2022 occurred during summer and involved a glaucous gull (*Larus hyperboreus*) and great skuas (*Stercorarius skua*) on Svalbard and Jan Mayen, representing the first detections of HPAIVs in the High Arctic. Three HPAIV genotypes were identified: EA-2020-C (H5N1), EA-2021-AB (H5N1), and EA-2021-I (H5N5). In 2023, HPAIVs were detected in a broader range of bird species, and retrospectively in an Atlantic walrus reported by another research group (*Odobenus rosmarus rosmarus*). Genotypes identified in 2023 were EA-2020-C (H5N1), EA-2021-I (H5N5), and EA-2022-BB (H5N1). No cases were reported in 2024. In 2025, EA-2021-I (H5N5) was detected in Arctic foxes (*Vulpes lagopus*) on Svalbard, without preceding detections in wild birds. The foxes exhibited neurological symptoms, and necropsy of one individual revealed the presence of feathers in its stomach. All sequenced viruses from the Arctic foxes uniquely carried the combination of PB2-E627K and PB1-H115Q, which is associated with mammalian adaptation.

The detection of multiple genotypes indicates repeated and independent introductions of HPAIVs into these regions. The co-circulation of genetically distinct virus strains in areas of high bird density further suggests that Arctic breeding grounds may facilitate local viral amplification, reassortment, and subsequent dissemination along migratory flyways, including transcontinental spread.

**Data summary:** The authors confirm all supporting data, code and protocols have been provided within the article or through supplementary data files. Influenza A whole genome sequences generated through this study are available under the GISAID accession numbers found in Table 1. All genome sequences and associated metadata supporting the findings of this study can be accessed through the persistent digital object identifier https://doi.org/10.55876/gis8.260211rq.

Table 1.
Detections of highly pathogenic avian influenza virus (HPAIV) in wild birds and mammals sampled in the Svalbard Archipelago and on Jan Mayen from January 1^st^, 2022, to August 31^st^, 2025.

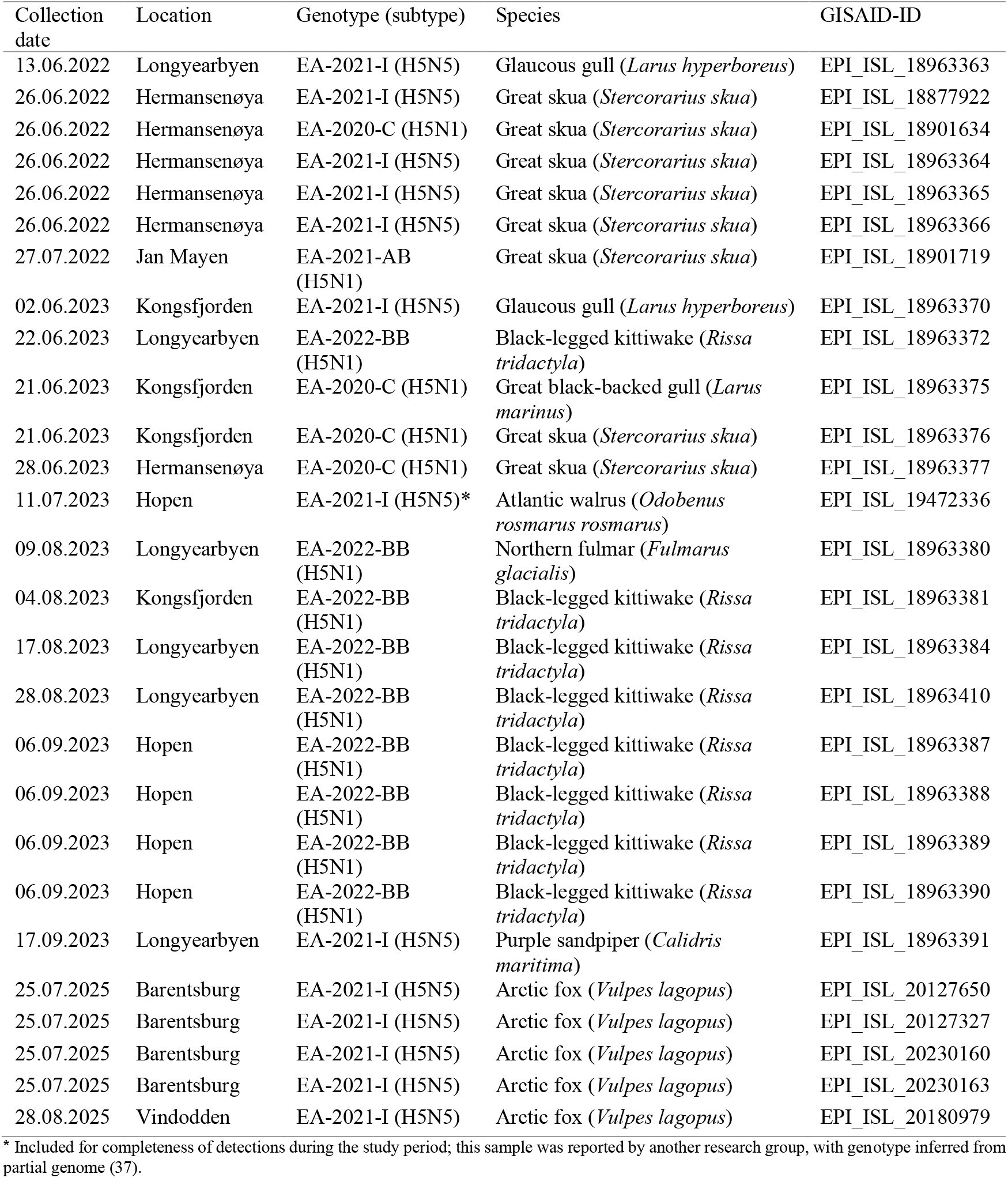

## Introduction

Since autumn 2020, highly pathogenic avian influenza viruses (HPAIVs) of H5Nx subtypes belonging to clade 2.3.4.4b of the A/goose/Guangdong/1/96 (Gs/Gd) lineage have caused substantial mortality in wild and domestic birds globally, with increasing spillover to mammals (1). The 2021–2022 epizootic wave marked a turning point, with rapid geographic expansion across Europe, Asia, Africa, and North America (2-6). HPAI H5Nx viruses became established in wild bird populations, suggesting enzootic circulation across several continents (7, 8). In 2022 and in 2023, cases were also confirmed in South America and the Antarctic region (9). During the summers of 2022 and 2023, HPAIV caused mass-mortality in colonies of breeding seabirds (10, 11).

HPAIVs possess high reassortment potential, and gene segment exchange with regionally circulating low pathogenic avian influenza viruses (LPAIVs) has generated a diverse pool of H5 Eurasian (EA) genotypes with distinct phenotypic characteristics (7, 12). The host range has also expanded considerably in recent years, encompassing a wide variety of mammalian species in addition to avian hosts. Adaptive mutations may enhance viral fitness in novel hosts, thereby increasing the probability of successful cross-species transmission (13). In mainland Norway, detections have largely reflected the pattern of genotypes predominating in Europe, with multiple H5Nx genotypes identified since 2020 (14-16). In contrast to the sporadic detections of other genotypes, EA-2021-I (H5N5) has been detected annually in Norway since 2022, suggesting continuous presence (17, 18).

In 2022, HPAIVs were detected for the first time in the Svalbard Archipelago and on the island of Jan Mayen (17), representing the first documented occurrence of HPAIV in the High Arctic. These remote regions, sparsely populated by humans, are ecologically significant breeding and feeding grounds for migratory birds. Each summer, millions of birds gather in dense colonies following migration from overwintering areas across the Atlantic and Arctic Oceans. Birds from diverse geographic origins may acquire infections through interactions with resident wild birds and domestic birds along migratory routes, facilitating long-distance dissemination of pathogens, including avian influenza viruses. Consequently, the High Arctic may function as a convergence zone for HPAIV exchange, similar to what has been proposed for Iceland (19, 20).

Svalbard is an Arctic archipelago located between the productive Greenland and Barents Seas. It hosts large numbers of birds, particularly seabirds (21-23). An estimated 3 million pairs of seabirds from approximately 18 species breed on the archipelago, with the little auk (*Alle alle)*, black-legged kittiwake (*Rissa tridactyla)*, northern fulmar (*Fulmarus glacialis)*, and common and Brünnich’s guillemots (*Uria aalge* and *U. lomvia)* being the most numerous (22, 24, 25). A few terrestrial migratory bird species also breed on Svalbard, with the barnacle goose (*Branta leucopsis)*, brent goose (*Branta bernicla)* and pink-footed goose (*Anser brachyrhynchus*) being the most abundant (26). Several waders are also common, of which the purple sandpiper (*Calidris maritima)* is the most prevalent (24). With the exception of the Svalbard rock ptarmigan (*Lagopus muta hyperborea*), all bird species are long-distance migrants that leave the archipelago during the non-breeding season to overwintering areas as far away as Antarctica, Africa, continental Europe, and the American coasts (27-29). Svalbard also hosts a diverse and abundant mammalian fauna. The terrestrial species comprise only the Arctic fox (*Vulpes lagopus)*, Svalbard reindeer (*Rangifer tarandus platyrhynchus*), and a small population of introduced sibling voles (*Microtus levis*) (30). In contrast, the diversity of marine mammals is much higher, including species such as polar bear (*Ursus maritimus*), Atlantic walrus (*Odobenus rosmarus rosmarus*) and multiple phocid seal species and cetaceans (30, 31).

Jan Mayen is a remote Arctic volcanic island located between the Greenland and Norwegian Seas. The island is 54 km long and covers 376 km^2^ (32). Its isolated location within a highly productive marine environment supports large seabird populations (33). The island is one of few sites where all six Atlantic auk species, the razorbill (*Alca torda*), common and Brünnich’s guillemot, black guillemot (*Cepphus grylle*), little auk and Atlantic puffin (*Fratercula arctica*), breed (34). In total, 18 seabird species nest on Jan Mayen distributed across 22 colonies comprising more than 300,000 breeding pairs (35). The terrestrial bird community is limited to a few species of waders and passerines, with the snow bunting (*Plectrophenax nivalis*) being the most common (32). All bird species breeding on the island are long-distance migrants that leave during the non-breeding season, and most seabirds are believed to overwinter in the northern Atlantic Ocean (35). Jan Mayen also serves as a stopover site for populations of geese and waders breeding in Greenland. The Arctic fox once bred on the island but went extinct following intensive hunting in the early 20th century (32).

In this study, we describe cases of HPAI in wild birds and mammals in the Svalbard Archipelago and on Jan Mayen between January 2022 and August 2025 to enhance understanding of HPAIV occurrence and transmission in the High Arctic ecosystem. We characterise the genomic diversity of detected viruses, focusing on phylogenetic relationships and mutations associated with mammalian adaptation, pathogenicity, and antiviral resistance.

## Materials & Methods

### Sampling

Passive surveillance for avian influenza in the Svalbard Archipelago and on Jan Mayen, defined as the sampling of dead or diseased wild birds or mammals, was conducted continuously throughout the study period (January 1^st^, 2022 – August 31^st^, 2025) whenever carcasses or symptomatic animals were encountered, reported, and could be sampled. The Governor of Svalbard, acting as the highest representative of the Norwegian government and the local authority on the archipelago, has issued clear guidelines urging residents, tourists, and researchers to report observations of sick or deceased wild birds and mammals (36, 37). The Governor’s nature management officers and field researchers collected samples and retrieved carcasses. All collected material, including swab samples, tissue samples, and/or carcasses, was submitted directly to the Norwegian Veterinary Institute for diagnostic examination and further analysis, following instructions from the Norwegian Food Safety Authority. The number of animals sampled per species through passive surveillance on Svalbard is listed in Supplementary Table 1. Additionally, samples from one Atlantic walrus were collected and analysed as described in Postel *et al*., 2025 (38).

Active surveillance for avian influenza was conducted on Svalbard in 2021 and 2022. No active surveillance was carried out on Jan Mayen during the study period (January 1^st^, 2022 – August 31^st^, 2025). The surveillance involved collecting cloacal and oropharyngeal swabs in transport medium from apparently healthy wild birds and was performed during the breeding season at selected colonies in connection with fieldwork such as bird ringing and population monitoring. In 2021, prior to the study period, a total of 98 barnacle geese were sampled on Svalbard, and no influenza A virus was detected (data not shown). In 2022, active surveillance primarily focused on waterfowl and seabird species (Supplementary Table 1). Samples collected or analysed outside the study period are not reported.

### Nucleic acid extraction and influenza A virus detection

The samples were analysed at the national reference laboratory for avian influenza at the Norwegian Veterinary Institute. Sample processing, influenza A virus screening by real-time reverse transcriptase PCR (rRT-PCR) targeting the *M* gene, and subtyping (H5, N1 and N5 assays) were performed as previously described (15, 39). Briefly, subtyping and pathotyping was performed using haemagglutinin (HA) and neuraminidase (NA) subtype- and HPAI H5 clade 2.3.4.4b specific rRT-PCR assays (40-42).

### Whole genome sequencing

For all samples, that tested positive for influenza A virus *M* gene by PCR, the EURL protocol based on methodology by Zhou *et al*. in 2009 (43), was followed to amplify the complete genomes (43). In brief, RNA extracted for virus identification was used as template for the reverse transcription-polymerase chain reaction (RT-PCR) amplification. This RT-PCR employed the SuperScript^TM^ III One-Step RT-PCR System with Platinum Taq HF DNA polymerase (Thermo Fisher Scientific) and the following primers: MBTUni-12-DEG-new (5’ – GCGTGATCAGCRAAAGCAGG – 3’) and MBTUni-13 (5’ – ACGCGTGATCAGTAGAAACAAGG – 3’), which are complementary to the conserved ends of the eight influenza A virus genome segments. The resulting RT-PCR products were purified using AMPure clean-up beads (Beckman Coulter) and analysed and quantified using a TapeStation (Agilent) automated capillary electrophoresis instrument. Two hundred nanograms DNA from each sample was used for Illumina DNA Prep (Illumina). Subsequently, the libraries were sequenced on the Illumina MiSeq or NextSeq systems.

### Genome assembly and sequence analyses

Most of the viral consensus sequences from the detections in the High Arctic were generated through reference-based mapping implemented in InsaFlu (44) using references as described in Bøe *et al*. (15). The HPAIV genomes of the five Arctic foxes were assembled using the Iterative Refinement Meta Assembler (IRMA) (45) v1.2.0. The viral consensus sequences generated in this study have been deposited in GISAID’s EpiFlu database (46). Genotyping was conducted following the method described by Fusaro *et al*. (7), using MEGA X v10.1.8 (47) by comparison with EA reference sequences provided by the EURL. All viral sequences were screened for mutations associated with mammalian adaptation, using FluMut GUI 3.1.1, FluMut 0.6.3, FluMutDB (48).

Similarity searches of all influenza A virus segments were performed separately using nucleotide sequences in GISAID EpiFlu’s BLAST tool (46). The analyses were based on metadata associated with sequences of subtype H5N1 available in GISAID up to November 10, 2024, or sequences of subtype H5N5 available on September 30^th^, 2025. The 50 most homologous nucleotide sequences were retrieved from each BLAST search. In addition, all sequences from Norwegian detections of the same subtype were included, as well as reference strains for each genotype and gene segment. Duplicate or incomplete sequences were removed manually. Spatiotemporally redundant sequences (originating from the same time and location) were removed manually to maintain HA phylogenies of approximately 50 representative sequences. Accession numbers for all included data are listed in Supplementary Table 2. Evolutionary analyses were conducted using MEGA X v10.1.8 (47) employing the MUSCLE algorithm with default parameters and bootstrap 1000 for nodal support. Evolutionary history was inferred by using the Maximum Likelihood method and Tamura-Nei model (49). The trees with the highest log likelihoods are shown. The trees are drawn to scale, with branch lengths measured in the number of substitutions per site. Some branches were compressed for easier visualisation.

### Epidemiological data and analysis

Submitted carcasses and samples, and their test results, were registered in the Norwegian Veterinary Institute’s laboratory management system (PJS version 21.1.25.0). R v4.5.1 (“Great Square Root”) (50) was used to retrieve, clean, interpret, aggregate and merge data with geocoordinates. The giscoR package (51) was used to download geospatial datasets, and the sf (52) and tmap packages for production of maps (53). Information regarding the HPAIV-positive Atlantic walrus was included from Postel *et al*., 2025 (38).

## Results

### Field observations and virus detections

Between January 1^st^, 2022, and August 31^st^, 2025, a total of 325 wild birds and 17 mammals from Svalbard and Jan Mayen (Figure 1A) were tested for avian influenza (Supplementary Table 1). Of these, 21 birds from six species and five mammals from one species tested positive for HPAIV H5Nx clade 2.3.4.4b by RT-PCR (Table 1). In addition, one Atlantic walrus tested positive for HPAIV as reported by Postel *et al*., 2025 (38).

**Figure 1.**
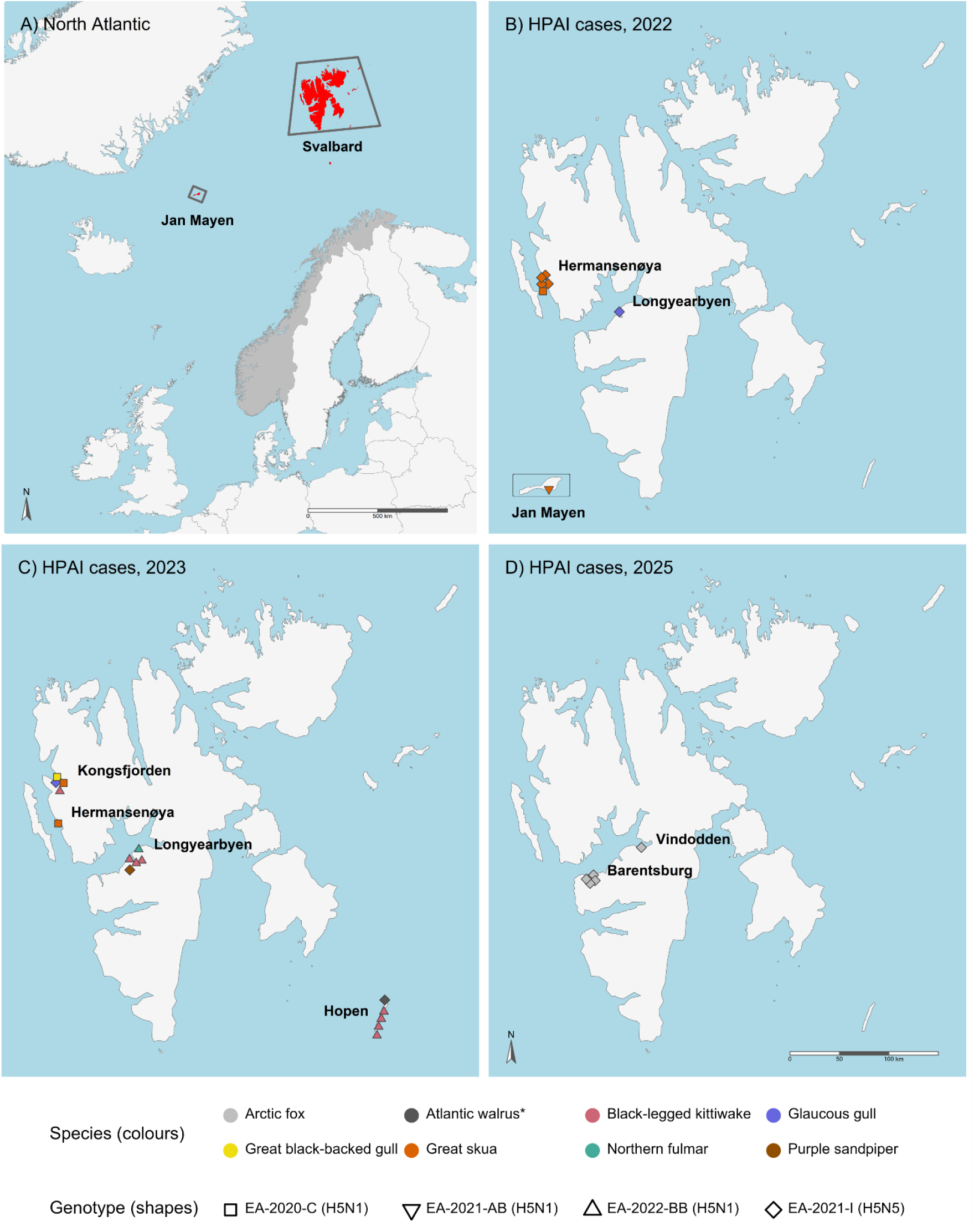
Maps of Svalbard and Jan Mayen (A) showing the locations of wild birds and mammals that tested positive for highly pathogenic avian influenza viruses (HPAIVs) in 2022 (B), 2023 (C), and 2025 (D). *Based on partial genome (38).

In 2022, HPAIV was detected in seven seabirds from two species in Svalbard and Jan Mayen. In June, EA-2021-I (H5N5) was detected in a glaucous gull (*Larus hyperboreus*) found dead by a dock near the airport in Longyearbyen, representing the first confirmed case of HPAIV in the Svalbard Archipelago and the High Arctic (Table 1, Figure 1B). Later the same month, HPAIV was detected in a breeding colony of great skuas (*Stercorarius skua*) on Hermansenøya, a small island located approximately 80 km northwest of Longyearbyen (Figure 1B). During a one-day survey, 27 dead great skuas were recorded on the island, indicating substantial mortality within the colony (Figure 2A). Of five individuals sampled, four tested positive for EA-2021-I (H5N5) and one for EA-2020-C (H5N1) (Table 1). Although barnacle geese, common eiders (*Somateria mollissima*), and glaucous gulls also breed on Hermansenøya, no unusual mortality was observed among these species during fieldwork at the time when great skua deaths were recorded. In July 2022, a great skua found dead on Jan Mayen tested positive for HPAIV EA-2021-AB (H5N1) (Table 1), documenting the presence of a third HPAIV H5 genotype in the High Arctic during the 2022 breeding season.

**Figure 2.**
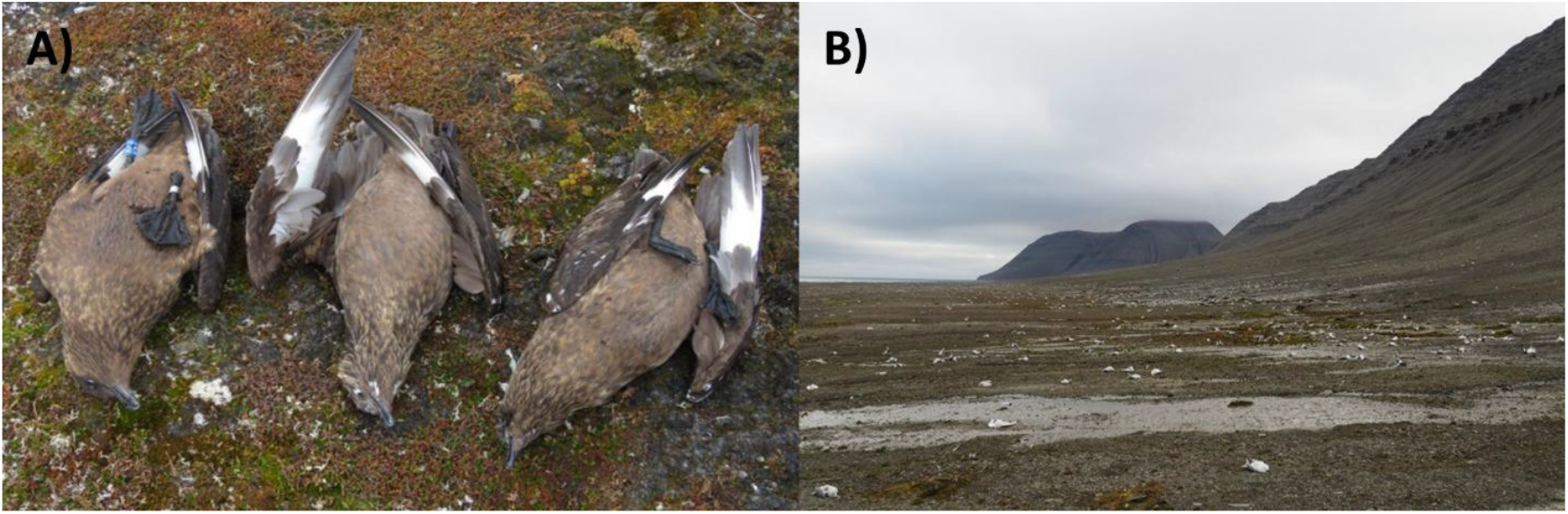
(A) Dead great skuas (*Stercorarius skua*) on Hermansenøya during the 2022 breeding season. Of five sampled individuals, four tested positive for the HPAIV genotype EA-2021-I (H5N5) and one for EA-2020-C (H5N1). (B) High mortality among black-legged kittiwakes (*Rissa tridactyla*) on Hopen in early September 2023. All sampled individuals tested positive for the HPAIV genotype EA-2022-BB (H5N1). Photos: Børge Moe (A) and Håvard Hansen (B).

In 2023, 14 wild birds and one marine mammal sampled across multiple locations in the Svalbard Archipelago tested positive for HPAIV by RT-PCR (Figure 1C). Samples were collected between June and September 2023 from multiple bird species (Table 1). The black-legged kittiwake was the most frequently affected species, accounting for eight positive cases. Two great skuas from separate locations (Kongsfjorden and Hermansenøya) tested positive, while single detections were made in glaucous gull, great black-backed gull (*Larus marinus*), northern fulmar, and purple sandpiper. Multiple EA H5 genotypes were identified among the detections.

The first detection of HPAIV in 2023 occurred in June, when EA-2021-I (H5N5) was identified in a glaucous gull found dead in Kongsfjorden (Figure 1C). The carcass was fresh at the time of sampling, with visible brood patches. Later that month, EA-2022-BB (H5N1) was detected in a black-legged kittiwake, while a great black-backed gull and two great skuas tested positive for EA-2020-C (H5N1). All four birds were found dead within the broader region of western Spitsbergen. The black-legged kittiwake was detected at Hotellneset near Longyearbyen, while the other birds were found in Kongsfjorden and on Hermansenøya.

In August and September 2023, high mortality was reported among seabirds on Svalbard (E. Rønning, the Governor of Svalbard, pers. comm.). The mortality was likely associated with EA-2022-BB (H5N1), which was detected in a northern fulmar found dead in Longyearbyen, and in seven black-legged kittiwakes from multiple locations (Table 1). The first positive kittiwake was a chick found dead in its nest in Kongsfjorden in the beginning of August. Two additional kittiwakes found in Longyearbyen later that month tested positive, and four individuals tested positive during a mass-mortality event on Hopen in early September (Figure 2B). The last detection of HPAIV in the High Arctic in 2023 was in a purple sandpiper found in Longyearbyen in September, which tested positive for the EA-2021-I (H5N5) genotype.

Retrospective testing performed by Postel *et al*. confirmed HPAIV H5N5 in an Atlantic walrus found dead on Hopen and sampled during summer 2023 (38). Reports of approximately 10– 20 dead walruses were received from islands in southern and western parts of the Svalbard Archipelago. Most carcasses were lost to the tide, and only one walrus on Hopen was available for sampling.

In 2024, HPAIV was not detected in wild birds or mammals in Svalbard or Jan Mayen. However, in July 2025, four Arctic fox cubs exhibiting neurological symptoms nearby the Russian settlement of Barentsburg, Svalbard, tested positive for EA-2021-I (H5N5) (Table 1, Figure 1D). The foxes were submitted to the Norwegian Veterinary Institute with a primary suspicion of rabies, due to the clinical signs and two previous confirmed rabies cases in Arctic foxes in Svalbard in 2025. All four foxes tested negative for rabies virus. No cases of HPAIV had been detected in wild birds in Svalbard in 2025 prior to this event. Necropsy revealed large quantities of feathers in the stomach of one of the foxes. In late August, a decomposed carcass of another Arctic fox cub was found at Vindodden, east of Longyearbyen, and submitted to the Norwegian Veterinary Institute with suspicion of rabies. This fox also tested negative for rabies virus and subsequently positive for HPAIV EA-2021-I (H5N5).

### Phylogenetic analysis

A total of 21 complete viral genomes were generated from wild bird samples. From mammalian samples, five genomes were generated in this study, and one was retrieved from GISAID. The results of the phylogenetic analyses conducted to investigate their genetic relationships are presented below by genotype.

#### EA-2021-I (H5N5)

Twelve EA-2021-I (H5N5) viral genome sequences were obtained from wild bird and mammalian samples collected on Svalbard. Four of the 2022 sequences originated from great skuas sampled on Hermansenøya on June 26 (Table 1). Phylogenetic analysis of the HA gene segment showed that three of the great skua-derived viral sequences clustered with H5N5 viruses detected in gull species, such as herring gull (*Larus argentatus*), glaucous gull, and great black-backed gull, sampled on mainland Norway the same year (Figure 3). The first HPAIV detection on Svalbard, a glaucous gull sampled on June 13 in Longyearbyen, also belonged to this cluster. The viruses from mainland Norway were detected both prior to (May) and concurrently with (mid-June) the first confirmed case on Svalbard. Notably, a virus detected in a red-tailed hawk (*Buteo jamaicensis*) sampled on April 15, 2023, on Prince Edward Island, Canada, also grouped within this cluster and shared highly similar HA sequences. In contrast, the fourth HA sequence from a great skua sampled on Hermansenøya in 2022 did not cluster with the others. This divergence was also evident across the remaining gene segments (Supplementary Figure 1A).

**Figure 3.**
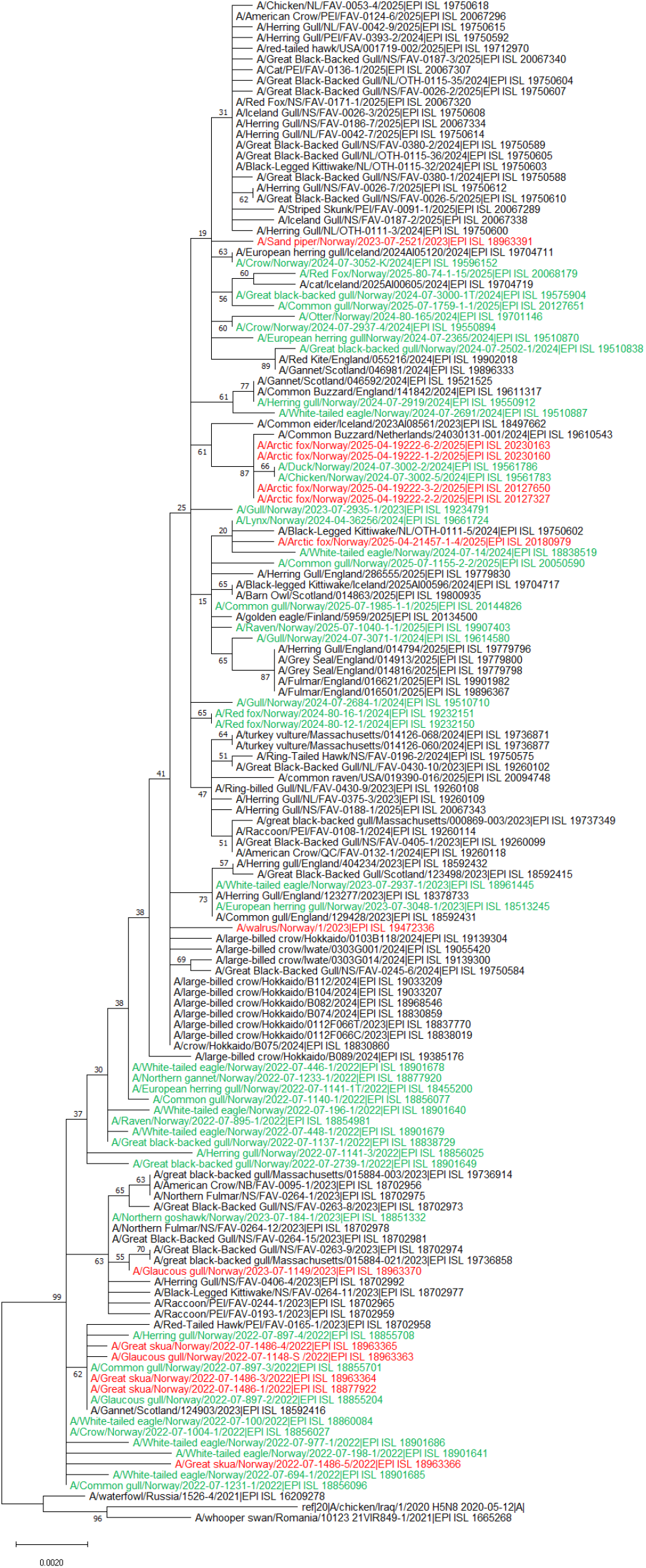
Maximum likelihood phylogenetic mid-point rooted tree of HA segments from highly pathogenic avian influenza viruses (HPAIVs) of genotype EA-2021-I (H5N5) detected in Svalbard, 2022-2025. Sequences from Svalbard are shown in red, and sequences from mainland Norway are shown in green.

The HA gene segment from the EA-2021-I (H5N5) virus detected in a glaucous gull in Kongsfjorden in early June 2023 showed relatedness to sequences from wild birds and mammals sampled in Canada in 2023, as well as to one sequence from a northern goshawk (*Accipiter gentilis*) sampled in Stavanger, southwestern Norway, in March 2023 (Figure 3). In contrast, the EA-2021-I (H5N5) virus detected in a purple sandpiper in Longyearbyen in mid-September 2023 was more closely related to a different group comprising sequences from viruses detected in wild birds and mammals in Norway, Northern Europe, and North America between 2024 and 2025 (Figure 3).

The HA sequence from the Atlantic walrus sampled by Postel *et al*. in summer 2023 showed the highest similarity to viruses detected in large-billed crows (*Corvus macrorhynchos*) in Hokkaido, Japan, the earliest of which was recorded in December 2023 (EPI_ISL_18837770). The viral genomes derived from Arctic foxes found in Barentsburg in July 2025 were more closely related to sequences from an outbreak in a backyard poultry holding on Frøya, Trøndelag, mainland Norway in November 2024, than to more recent Norwegian detections of EA-2021-I (H5N5) in wild birds and mammals (Figure 3, Supplementary Figure 1A). Finally, the HA sequence from the Arctic fox found in late August 2025 at Vindodden differed from those of the Barentsburg foxes and showed closer relatedness to sequences from other recent detections in Norway and around the Norwegian Sea.

#### EA-2020-C (H5N1)

The EA-2020-C (H5N1) virus sequence detected in a great skua on Hermansenøya in 2022 did not cluster with sequences of the same genotype identified from the same species and location during the subsequent summer (Figure 4A, Supplementary Figure 1B). The 2022 virus was positioned on a distinct branch of more European origin and showed highest similarity to viruses detected in great skuas in Scotland in 2021. It also shared high similarity to a partial genome (segments NP, NA, MP and NS) obtained from a white-tailed eagle sampled on mainland Norway in 2022. This branch also included viruses detected in geese, gulls and raptors sampled across Europe in 2021. In contrast, the sequences obtained from great skuas and a great black-backed gull, all three sampled within a single week in June 2023, formed a well-supported monophyletic cluster. This cluster grouped with sequences from scavenger species such as crows and raptors in Alaska, Eastern Russia and Hokkaido 2022 and 2023.

**Figure 4.**
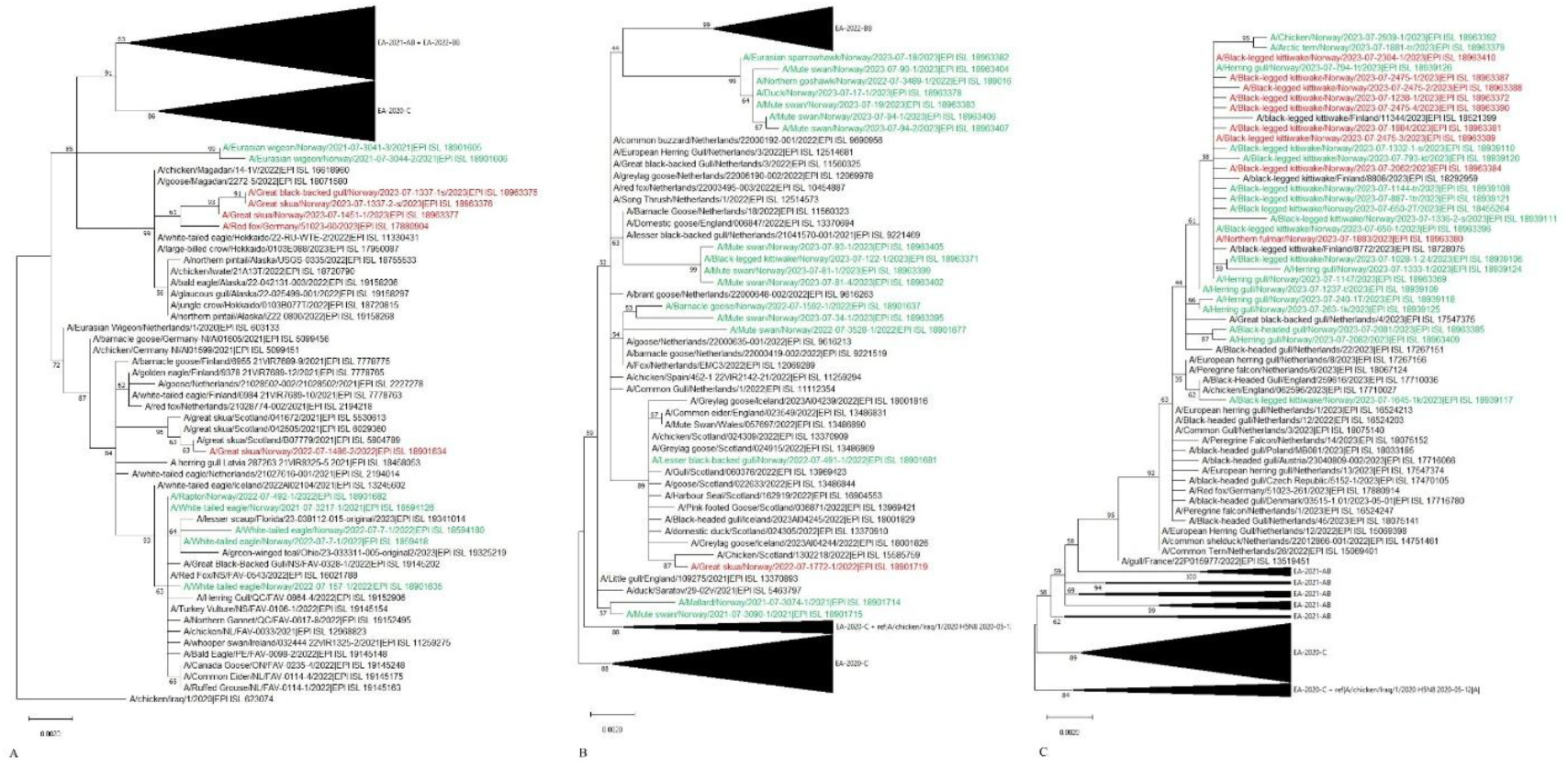
Maximum likelihood phylogenetic mid-point rooted tree of HA segments from highly pathogenic avian influenza viruses (HPAIVs) belonging to genotype EA-2020-C (A), genotype EA-2021-AB (B), and genotype EA-2022-BB (C) detected in Svalbard and Jan Mayen, 2022-2025. Sequences from Svalbard and Jan Mayen are shown in red, and sequences from mainland Norway are shown in green. The HA sequences obtained from birds on mainland Norway and belonging to other genotypes are compressed for simplicity.

#### EA-2021-AB (H5N1)

During the study period, EA-2021-AB (H5N1) was detected in only one wild bird from the Norwegian part of the High Arctic: a great skua sampled on Jan Mayen in late July 2022. In the phylogenetic analysis, the closest related HA sequence originated from a chicken in Scotland, sampled in October 2022 (Figure 4B). The virus detected in Jan Mayen showed lower genetic similarity to other EA-2021-AB viruses detected on mainland Norway, with one exception: a virus from a lesser black-backed gull (*Larus fuscus*) sampled in Bergen, in early May 2022. All gene segments of this virus clustered within the same phylogenetic group as the Jan Mayen detection (Figure 4B, Supplementary Figure 1C) and included viral sequences from seabirds sampled in Scotland, England, Wales, and Iceland, as well as Scottish poultry.

#### EA-2022-BB (H5N1)

Multiple detections of EA-2022-BB (H5N1) were made in Svalbard during August and September 2023, including one northern fulmar and several black-legged kittiwakes sampled in Longyearbyen, Kongsfjorden, and Hopen. These followed an earlier detection in June 2023, when a black-legged kittiwake was sampled in Longyearbyen. The gene segments from these birds clustered within the same broadly defined phylogenetic group comprising mostly isolates from the Norwegian mainland, and the June sequence showed no phylogenetic distinction from the later detections (Figure 4C, Supplementary Figure 1D). Genetically related viruses were also detected in three black-legged kittiwakes found inland in Finland (Inari and Kemijärvi) in July and September 2023.

### Mutation screening

The FluMut analysis showed that the viruses detected on Svalbard and Jan Mayen were mainly avian-adapted, but all contained several substitutions associated with mammalian adaptation (Supplementary Table 3). All viruses contained mutations in HA, among which S133A, S154N and T156A were present in the avian H5N1 viruses and S133A and S154 were detected in both the avian and mammalian H5N5 viruses. The H5N1 EA-2022-BB viruses also contained NP-Y52N and NA-S369I. The only mutation uniquely identified in all the mammalian H5N5 viruses was PB2-E627K. The viruses from the Arctic foxes also contained PB1-H115Q.

## Discussion

Current knowledge about the occurrence and distribution of avian influenza viruses (AIVs) in the High Arctic remains limited. Prior to the 2020/2021 epizootic wave, virus circulation in wild birds in this region was poorly documented. However, the detection of influenza A virus antibodies in clinically healthy gulls in Svalbard suggests that exposure to LPAIVs had occurred (54). In the present study, we show that Svalbard and Jan Mayen have experienced multiple independent introductions of HPAIVs in recent years. Our findings suggest that these introductions may have originated from both mainland Norway and the broader North Atlantic region. This indicates that avian breeding grounds in the High Arctic both influence and are influenced by virus transmission along multiple migratory routes involving diverse bird species.

Hermansenøya was a key site of virus detection in this study. Two distinct HPAIV genotypes, EA-2021-I (H5N5) and EA-2020-C (H5N1), were detected in the island’s great skua breeding colony in June 2022. The high bird density during the breeding season may facilitate efficient virus transmission, and co-circulation of different AIVs could allow viral reassortment in cases of co-infection. We found no evidence of genetic reassortment in our data, as all gene segments grouped consistently within their respective genotypes. Interestingly, among the EA-2021-I (H5N5) detections on Hermansenøya, one virus clustered separately from the others. This divergence may indicate a separate introduction event or local viral evolution within the colony. Colony counts on Hermansenøya indicated a marked decline in nesting activity among great skuas between 2021 and 2022, from 137 to 59 nests (a 57% reduction), coinciding with the HPAIV outbreak and suggesting a substantial impact on the breeding population (G. W. Gabrielsen, Norwegian Polar Institute, pers. comm.).

In 2023, genotype EA-2020-C (H5N1) re-emerged in great skuas in the breeding colony at Hermansenøya. Phylogenetic analysis revealed no clustering between the 2022 and 2023 sequences, indicating that the virus did not persist locally between seasons via overwintering in frozen carcasses or other environmental reservoirs. Instead, the sequence obtained in 2023 clustered with viruses detected in a great black-backed gull and another great skua from Kongsfjorden on Svalbard, all sampled in June 2023. This phylogenetic pattern strongly suggests that the 2023 detections reflect a new introduction of EA-2020-C (H5N1) into Svalbard rather than continued local circulation.

Most HPAIV-positive birds detected in Svalbard and Jan Mayen during 2022–2023 were seabirds. Gulls have been identified as key drivers of HPAI H5 dissemination, likely due to their long-distance movements and limited prior exposure and immunity to this subtype (55). Seabirds typically breed in dense colonies and frequently aggregate socially during both breeding and foraging, which may facilitate viral transmission. The risk of HPAIV introduction and spread within colonies is further increased by scavenging behaviour. Glaucous gulls, great black-backed gulls, and great skuas frequently feed on carcasses of birds and marine mammals, increasing their potential exposure to HPAIVs. Skua species have been proposed as key species for studying virus evolution and transmission in Antarctica (56), and our findings support a similar role in the High Arctic.

The EA-2022-BB (H5N1) genotype caused high mortality among gull species across Europe during the spring and summer of 2023, including a large outbreak in black-legged kittiwakes in Northern Norway (7, 39). Phylogenetic analysis of EA-2022-BB (H5N1) indicates that following introduction to Svalbard, the virus spread to multiple locations. The first detection was documented in June 2023 in a single kittiwake in Longyearbyen, with subsequent detections at several sites, including Hopen, in August and September (Figure 1B). This pattern indicates that local spread in the Svalbard Archipelago likely began in early summer and continued throughout the season. Multiple introductions from mainland Norway cannot be excluded. EA-2022-BB (H5N1) viruses from Svalbard and mainland Norway were highly genetically similar (Figure 3D), indicating that they were part of the same outbreak, which likely spread from Europe into western Norway before moving northwards (30). Although large-scale movements of adult birds are not expected during the breeding peak, movements of failed breeders and possibly non-breeders to prospect other colonies have been documented (57-59). Moreover, HPAIV infection may disrupt normal behaviour and alter movement patterns. The inland locations in Finland where EA-2022-BB (H5N1)-positive kittiwakes were detected in 2023 are unusual for this seabird species. Atypical movements potentially associated with HPAIV infection have been reported for other seabirds, such as northern gannets (*Morus bassanus*) (51, 52).

Prior to the study period, in September 2021, coinciding with HPAI detections in great skua colonies on islands off the shore of Scotland (60), a disease outbreak with high mortality and clinical signs consistent with HPAI was observed on Jan Mayen. Samples never reached the Norwegian Veterinary Institute, and the suspected infection was therefore not verified. The detection of EA-2021-AB (H5N1) in a single wild bird from Jan Mayen in 2022 suggests that this genotype was not widespread in the Norwegian High Arctic during the study period. However, undetected EA-2021-AB (H5N1) circulation on Svalbard cannot be ruled out. Phylogenetic analysis shows that the most similar virus in Norway was detected in a lesser black-backed gull from Bergen in the southern part of mainland Norway, with related sequences reported from Scotland, England, Wales, and Iceland. The island’s geographic location may expose it to different viral introduction pathways from mainland Europe and Greenland than those primarily influencing Svalbard. Subsequent detections indicate that Jan Mayen continues to receive occasional viral incursions. In September 2025, after the period covered by this study, EA-2021-I (H5N5) was detected in wild birds on Jan Mayen, specifically in glaucous gulls and in a lesser black-backed gull (EPI_ISL_20216223, data not shown).

The phylogenetic analysis suggests that at least six separate incursions of the EA-2021-I (H5N5) genotype occurred in Svalbard between 2022 and 2025, indicating repeated introductions rather than a single persistent lineage. Most of the viral genomes obtained from Svalbard were related to sequences from mainland Norway, suggesting that seabird populations along the Norwegian coast represent a recurrent source. The phylogenetic patterns also indicate transatlantic dissemination of EA-2021-I (H5N5), as a red-tailed hawk sampled in Canada in April 2023 carried a virus closely related to the 2022 Svalbard H5N5 viruses, highlighting the potential for long-distance spread via migratory birds. Transatlantic spread of HPAIVs has been documented previously (19, 20), and several seabirds common in Svalbard, including black-legged kittiwakes and great skuas, migrate to Canadian waters during the non-breeding season (61, 62), providing opportunities for viral exchange. Erdelyan *et al*. (63) further confirmed that the H5N5 virus from a glaucous gull sampled in Longyearbyen in June 2022 clustered phylogenetically with the virus detected in the red-tailed hawk in Canada (63).

EA-2021-I (H5N5) was also detected in Arctic mammals during the study period. Spillover of this genotype from birds to mammals has been increasingly reported in recent years, including a human case in the USA (63-65). In agreement with Postel *et al*. (38), our phylogenetic analysis confirmed that the HPAIV detected in the Atlantic walrus sampled on Svalbard in 2023 was genetically similar to strains later detected in wild birds in Japan, suggesting a broader Northern Hemisphere dissemination of this genotype. We detected EA-2021-I (H5N5) in Arctic foxes on Svalbard in two occasions in 2025, marking the first report of H5N5 in this species. Viruses from Barentsburg (July) and Vindodden (August) differed genetically, indicating sporadic and independent bird-to-mammal spillover events rather than mammal-to-mammal transmission. The phylogenetic relatedness between the virus detected in Arctic foxes in Barentsburg 2025 and a 2024 backyard poultry outbreak on mainland Norway raises questions regarding the mechanisms and pathways by which HPAIVs reach and potentially persist in the High Arctic. The temporal gap and the geographic distance between the detections, together with the lower similarity to more recent H5N5 viruses in wild birds and mammals in Norway, suggest that several H5N5 strains may have co-circulated, with some viruses persisting undetected for extended periods before re-emerging in new locations and hosts. Such persistence may be facilitated by virus maintenance in remote, sparsely inhabited regions and by circulation in subclinically infected or small bird species whose carcasses are rarely found.

The mutation analysis indicated that the viruses detected in both birds and mammals remained predominantly avian adapted. However, several mutations that may enhance binding to human-like influenza receptors were identified in the HA segments (66). In addition, the EA-2022-BB (H5N1) viruses harboured single amino acid substitutions in NP and NA that could increase their zoonotic potential (67). Notably, the PB2-E627K substitution, associated with increased replication efficiency in mammalian cells, was detected in the H5N5 viruses from mammals, supporting the notion that PB2-E627K is selected for during replication in mammalian hosts (68). In addition, the viruses from all the Arctic foxes contained PB1-H115Q. The combination of PB2-E627K and PB1-H115Q has previously been associated with aerosol transmission of H7N9 among guinea pigs, and has also been observed in recent H5N1 clade 2.3.2.1e strains from human cases in Cambodia and Vietnam (65, 69, 70).

All detections of AIVs in the Norwegian sector of the High Arctic to date have been made through passive surveillance, underscoring the importance of having operational systems and resources readily available to sample dead or diseased wild birds and mammals. Due to the remoteness of the region, the reported numbers of dead birds likely underestimate both the true number of cases and the geographic extent of the disease. Most samples from Svalbard and Jan Mayen were collected during the breeding season and in areas with known human activity, and our results therefore primarily reflect this period and these geographic locations. Fieldwork during the polar night (mid-November to late January) is minimal because of darkness and safety concerns, making mortality events difficult to detect. Combined with the absence of most migratory bird species, this likely explains the low number of HPAIV detections in winter.

In this study, no specific analyses were conducted to investigate potential introduction routes or sources of HPAIVs, such as time-resolved phylogenetics or Bayesian stochastic search variable selection. Consequently, our study provides only limited insight into the spatiotemporal dynamics and directionality of virus introductions to the Svalbard Archipelago and Jan Mayen, with only general patterns inferred from the phylogenetic trees. Future studies could address these limitations by integrating larger genomic datasets with complementary bioinformatic approaches to better resolve transmission pathways.

## Conclusions

Our findings demonstrate high genetic diversity and repeated introductions of HPAIVs into Svalbard and Jan Mayen. These results underscore the need for sustained and robust avian influenza surveillance in the High Arctic to detect emerging viral variants, improve understanding of transmission dynamics, and track the movement of viruses along migratory flyways, thereby strengthening preparedness in this region and along migratory routes.

## Author contributions

The project was conceived by CAB, SG, BG and KM. The experiments were carried out and data analysed by CAB, SG, RT, BG and JÅ. The manuscript was drafted by SG, CAB, JÅ and RT and edited and revised by all authors.

## Ethical approval

This study did not involve human participants, human material, or human data. Therefore, ethical approval from a human research ethics committee was not required.

## Ethical statement

All birds and mammals included in this study were found dead prior to sampling. The only exception were the Arctic foxes from Barentsburg, which were humanely euthanized by the Governor of Svalbard due to a primary suspicion of rabies, based on observed clinical signs and animal welfare considerations. All carcasses were sampled within the framework of national surveillance and monitoring programs. No animals were euthanized for the specific purposes of this study.

## Acknowledgements

We gratefully acknowledge all data contributors, i.e., the Authors and their Originating laboratories responsible for obtaining the specimens, and their Submitting laboratories for generating the genetic sequence and metadata and sharing via the GISAID Initiative, on which this research is based (see Supplementary Table 2 and EPI_SET_260211rq GISAID identifier). We wish to thank all field personnel, wildlife officers, and researchers involved in the collection of samples from wild birds and mammals. Their efforts were essential for the successful implementation of both passive and active surveillance in the Svalbard Archipelago and Jan Mayen. We sincerely thank the laboratory staff at the Norwegian Veterinary Institute for their excellent work in performing analyses and laboratory procedures.

