## Supplementary Figure 1 for "Multiple Introductions of Highly Pathogenic Avian Influenza Viruses into the High Arctic: Svalbard and Jan Mayen, 2022–2025"

### Supplementary Figure Legends

**Supplementary Figure 1A – Phylogeny EA-2021-I (H5N5)**
Maximum likelihood phylogenetic mid-point rooted trees of PB2/PB1/PA/NP/NA/M/NS segments (p. 1-7) from highly pathogenic avian influenza viruses (HPAIVs) of genotype EA-2021-I (H5N5) detected in Svalbard, 2022-2025. Sequences from Svalbard are shown in red, and sequences from mainland Norway are shown in green.

**Supplementary Figure 1B – Phylogeny EA-2020-C (H5N1)**
Maximum likelihood phylogenetic mid-point rooted trees of PB2/PB1/PA/NP/NA/M/NS segments (p. 1-7) from highly pathogenic avian influenza viruses (HPAIVs) belonging to genotype EA-2020-C detected in Svalbard, 2022-2023. Sequences from Svalbard are shown in red, and sequences from mainland Norway are shown in green. The sequences obtained from samples on mainland Norway and belonging to other genotypes are compressed for simplicity.

**Supplementary Figure 1C – Phylogeny EA-2021-AB (H5N1)**
Maximum likelihood phylogenetic mid-point rooted trees of PB2/PB1/PA/NP/NA/M/NS segments (p. 1-7) from highly pathogenic avian influenza viruses (HPAIVs) belonging to genotype EA-2021-AB detected in Svalbard, 2022-2023. Sequences from Jan Mayen are shown in red, and sequences from mainland Norway are shown in green. The sequences obtained from birds on mainland Norway and belonging to other genotypes are compressed for simplicity.

**Supplementary Figure 1D – Phylogeny EA-2022-BB (H5N1)**
Maximum likelihood phylogenetic mid-point rooted trees of PB2/PB1/PA/NP/NA/M/NS segments (p. 1-7) from highly pathogenic avian influenza viruses (HPAIVs) belonging to genotype EA-2022-BB detected in Svalbard 2022-2023. Sequences from Svalbard are shown in red, and sequences from mainland Norway are shown in green. The sequences obtained from birds on mainland Norway and belonging to other genotypes are compressed for simplicity.

### EA-2021-I, segment PB2

Supplementary Figure 1A

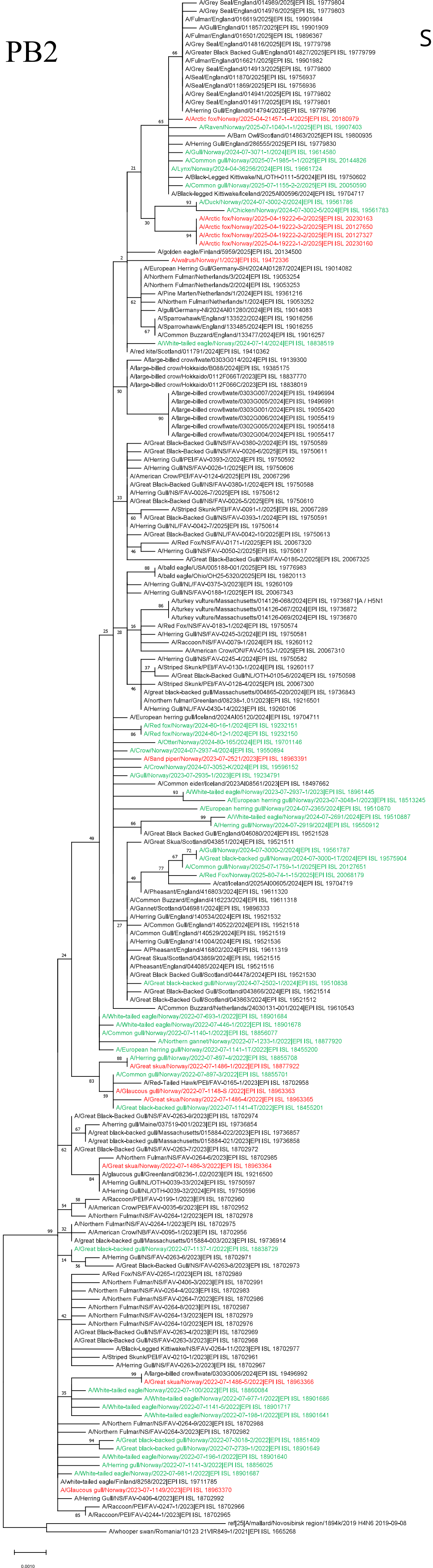

### EA-2021-I, segment PB1

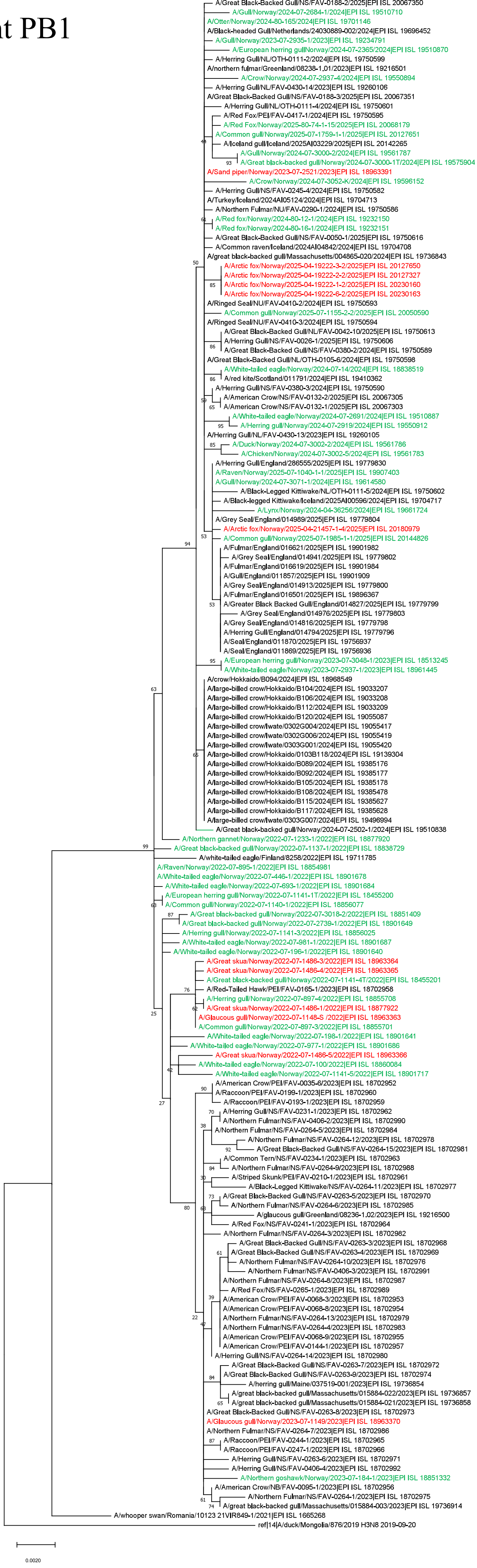

EA-2021-I, segment PA

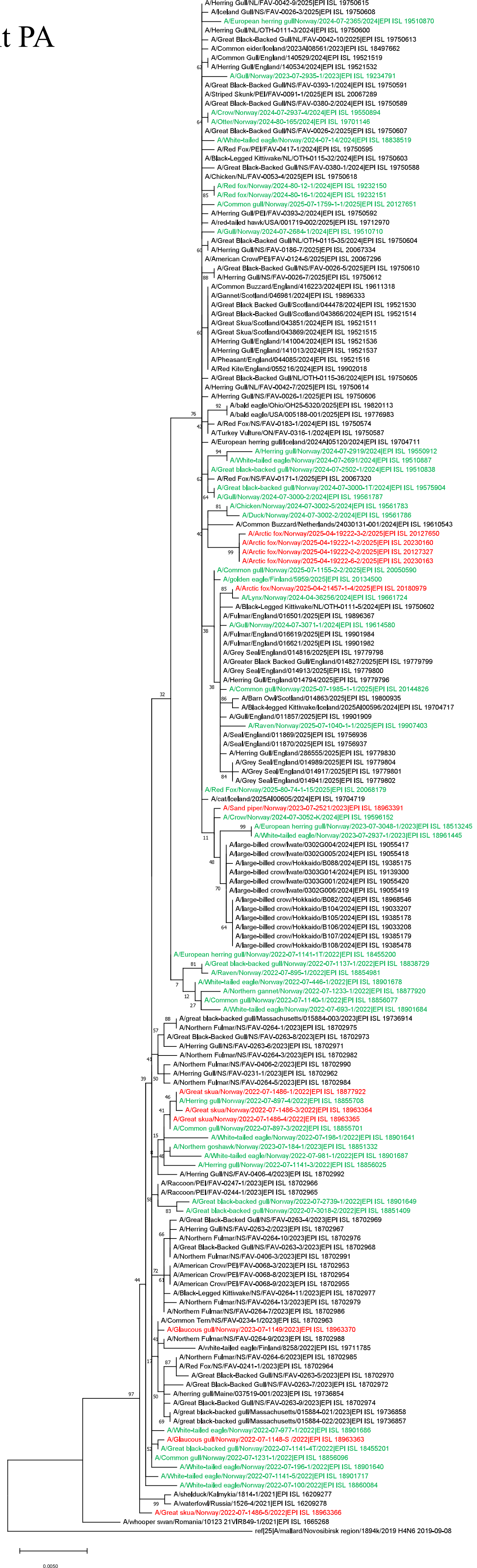

### EA-2021-I, segment NP

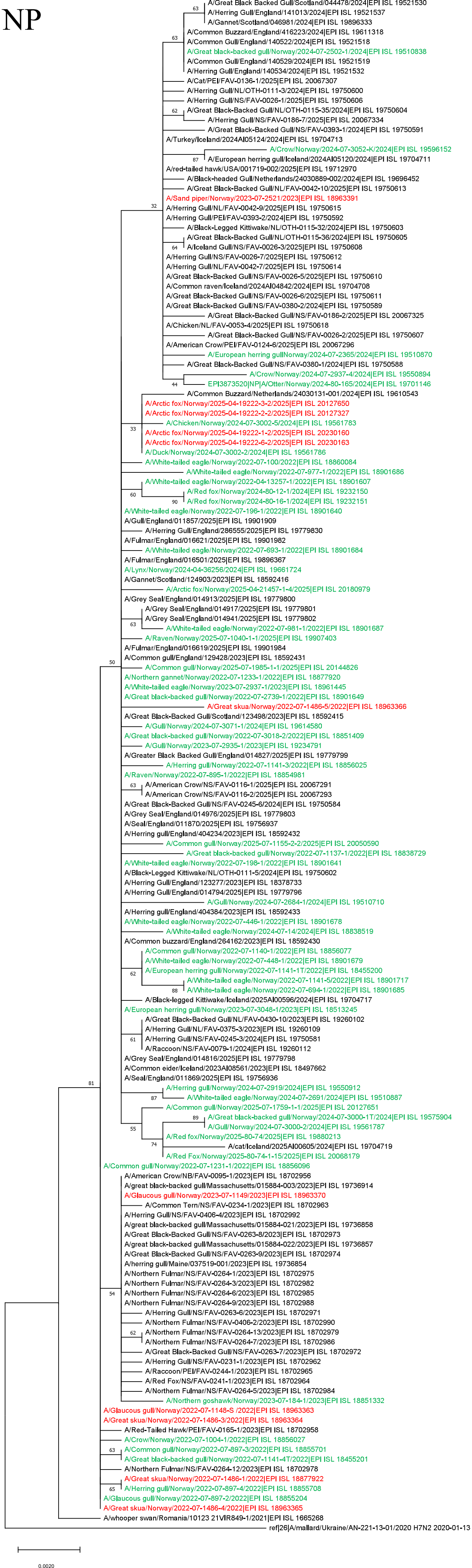

### EA-2021-I, segment NA

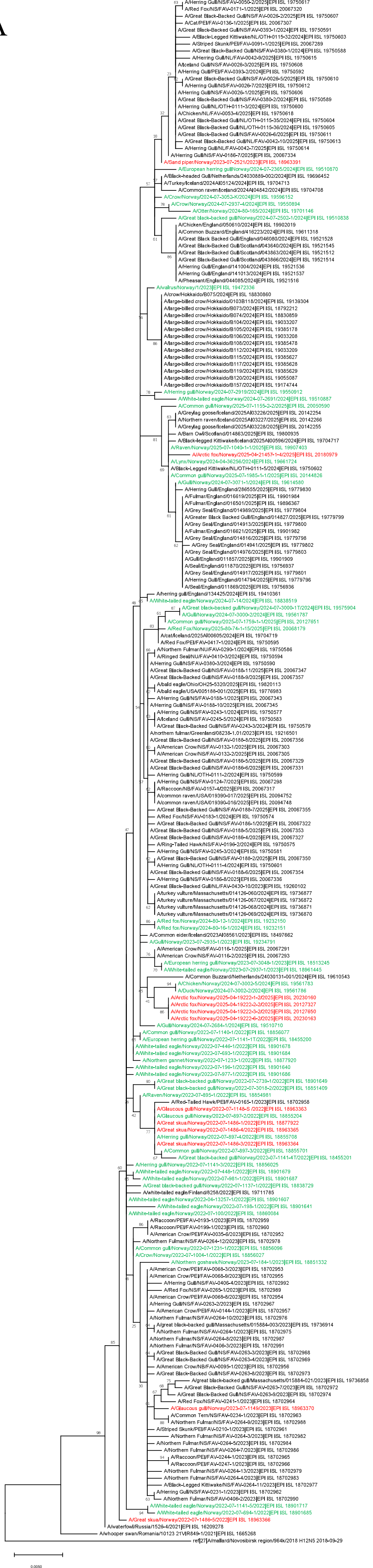

EA-2021-I, segment MP

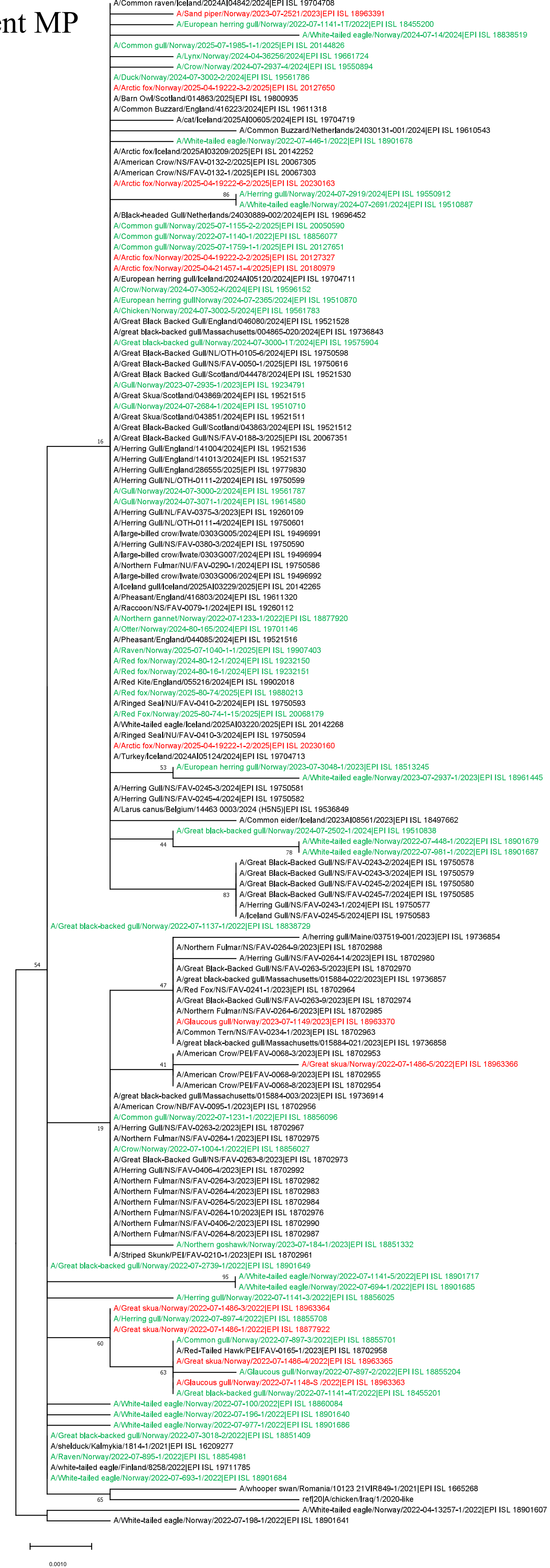

### EA-2021-I, segment NS

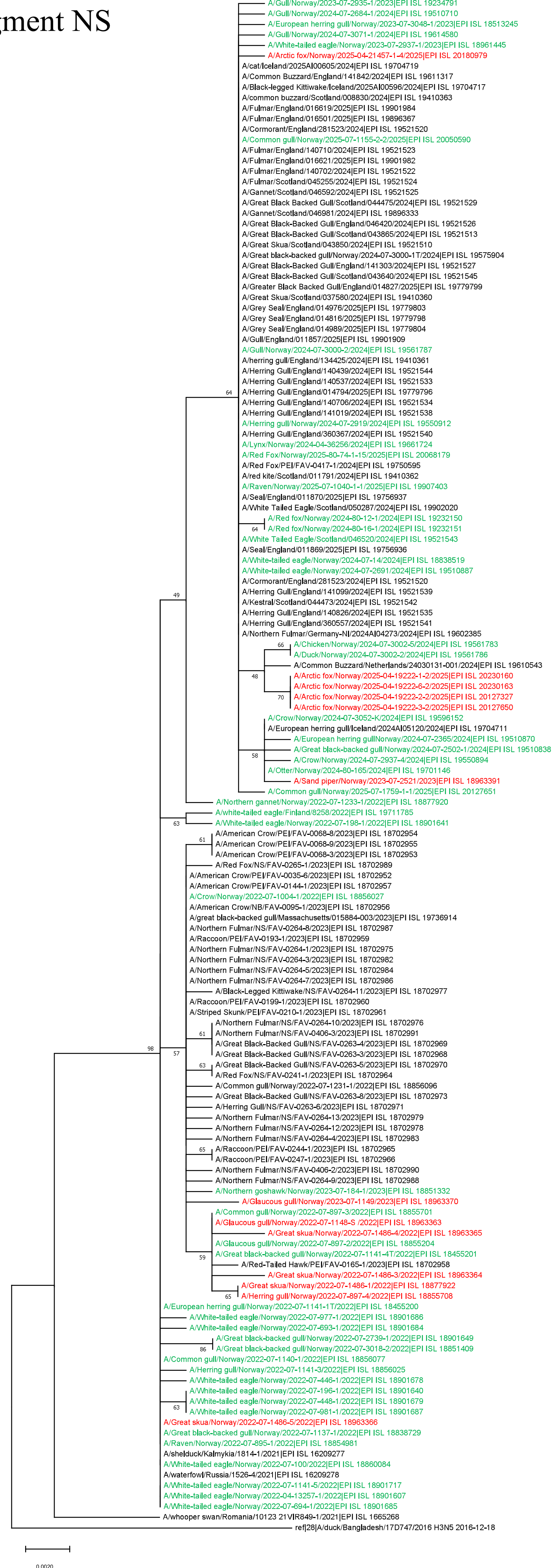

EA-2020-C, segment PB2

Supplementary Figure 1B

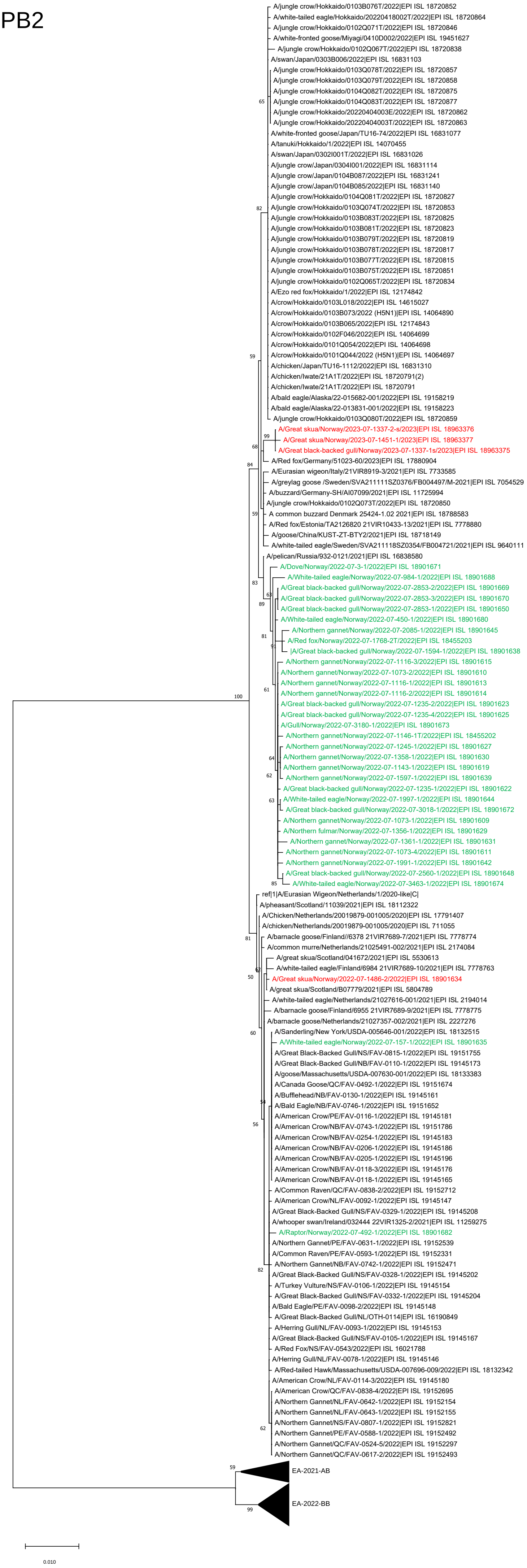

### EA-2020-C, segment PB1

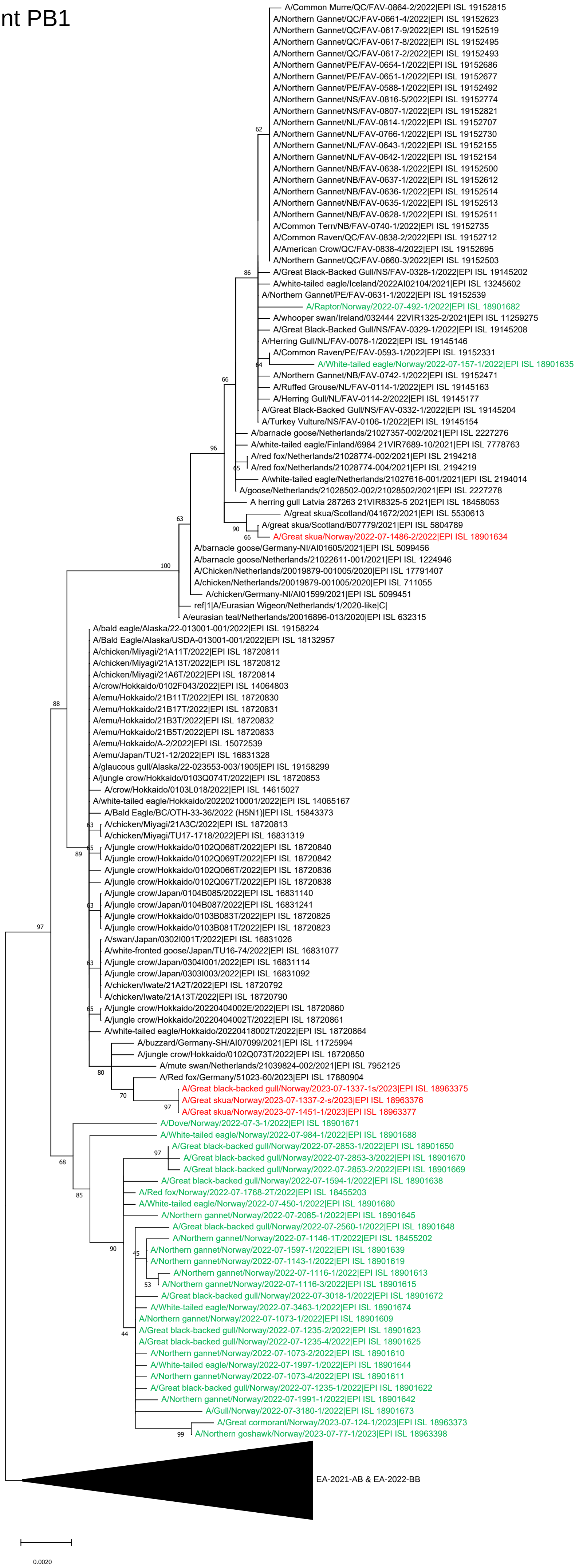

### EA-2020-C, segment PA

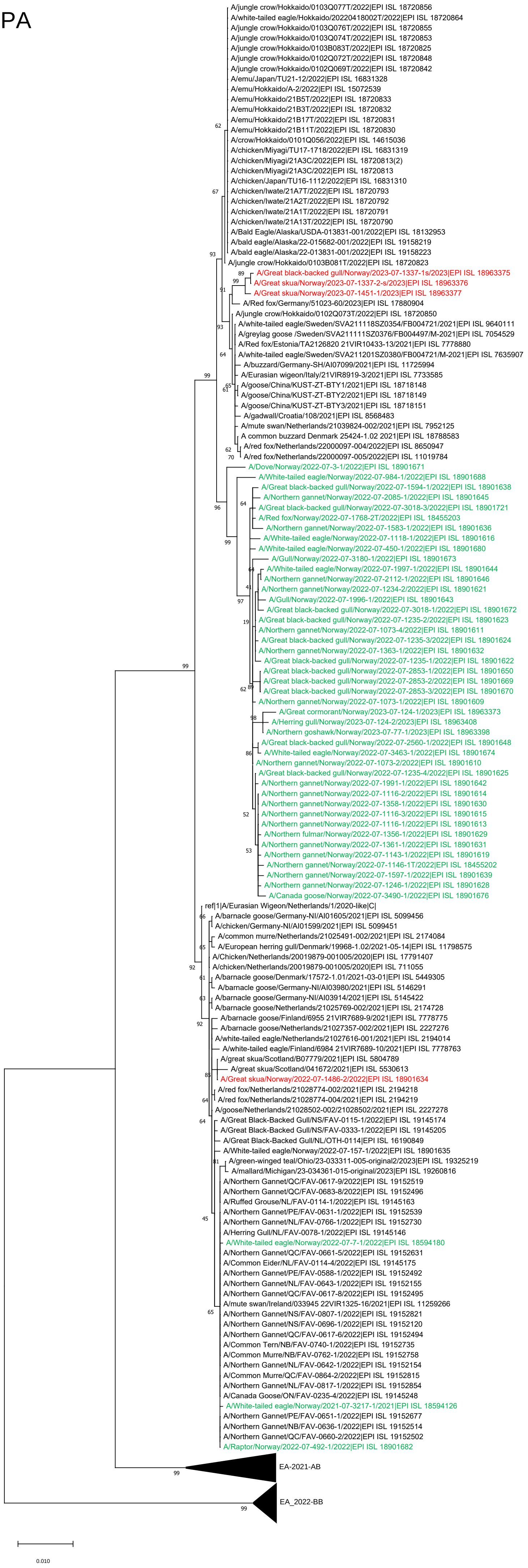

### EA-2020-C, segment NP

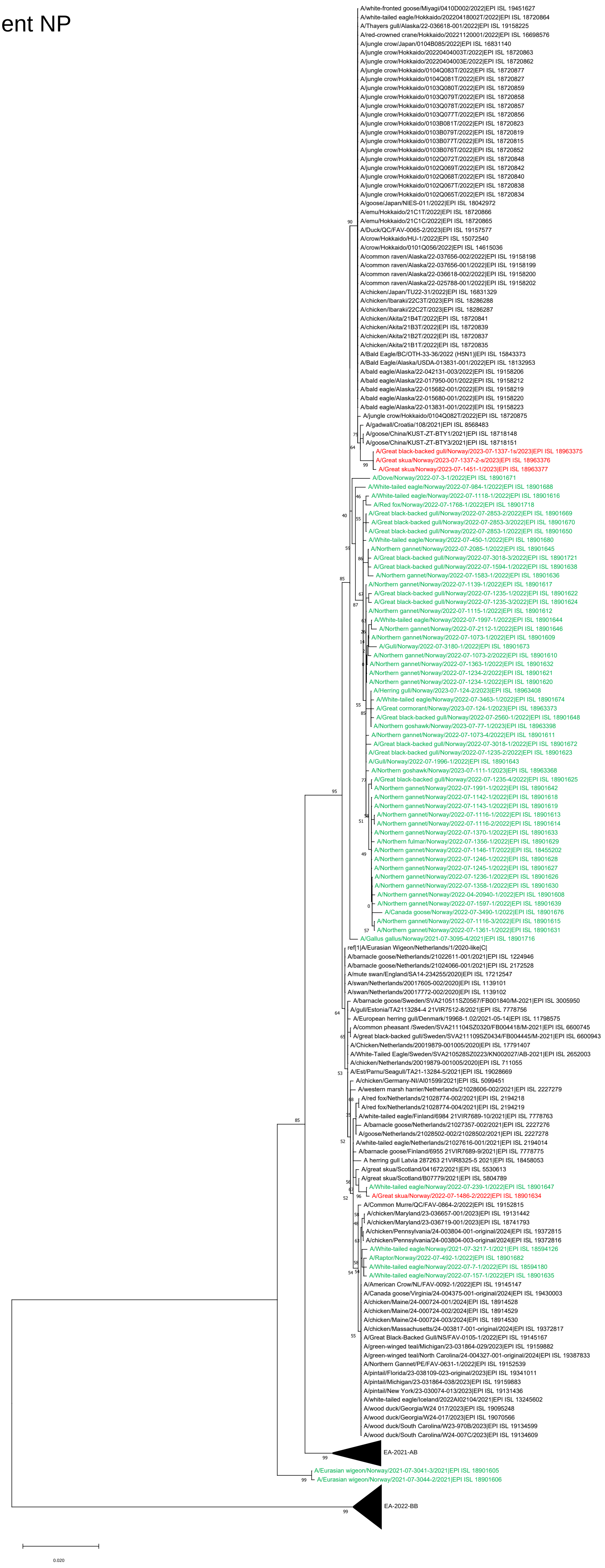

### EA-2020-C, segment NA

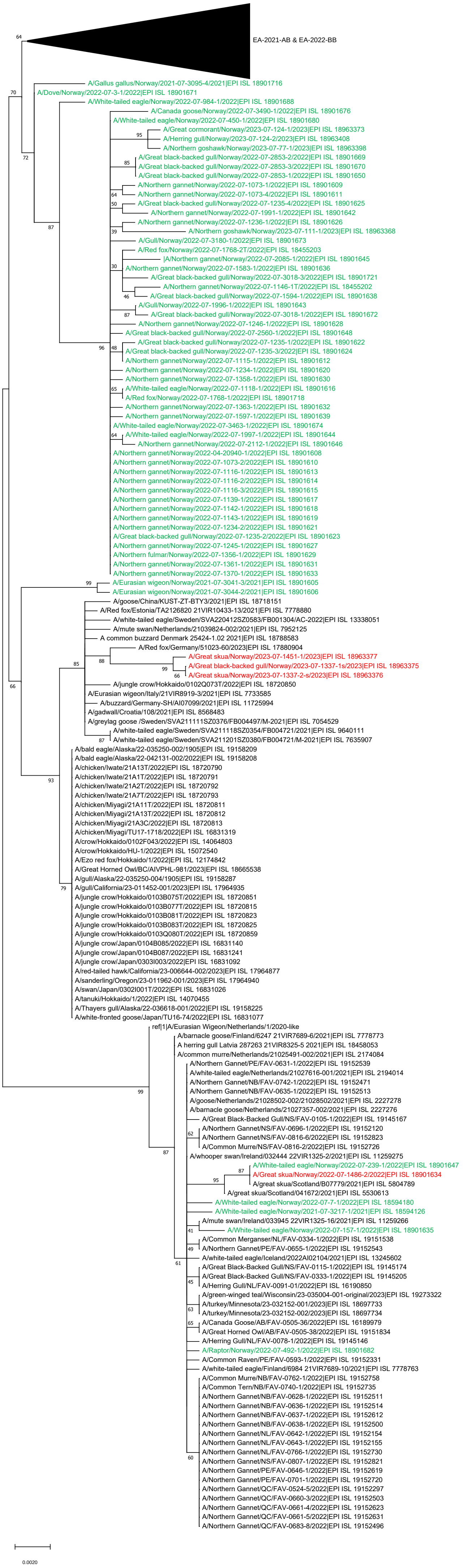

### EA-2020-C, segment MP

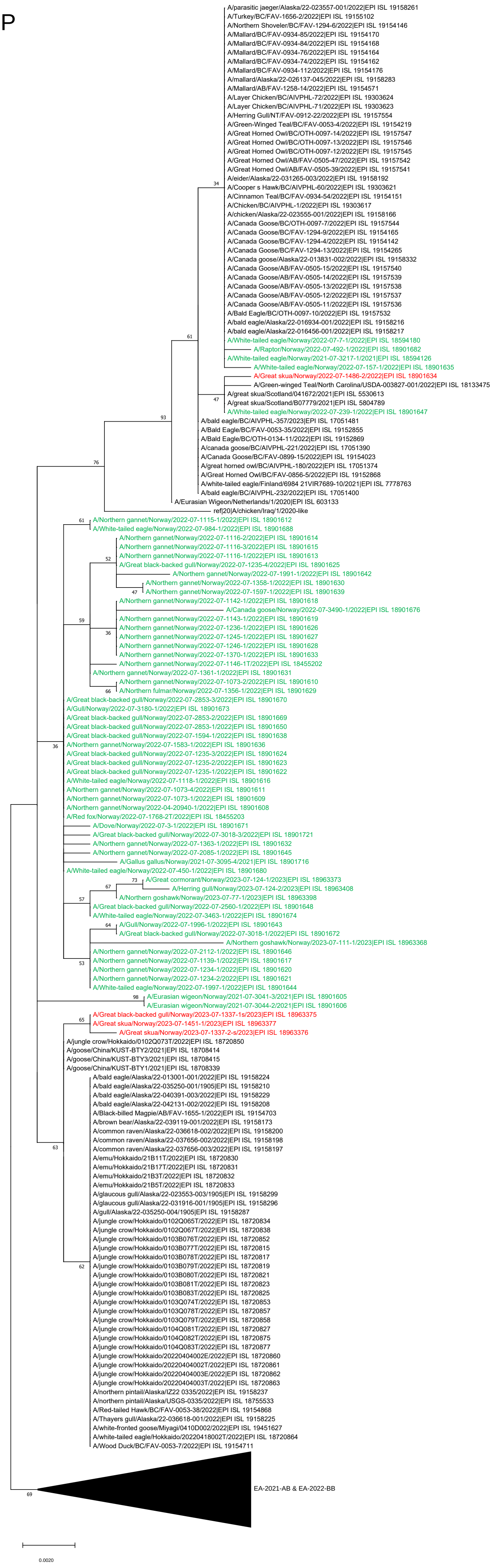

### EA-2020-C, segment NS

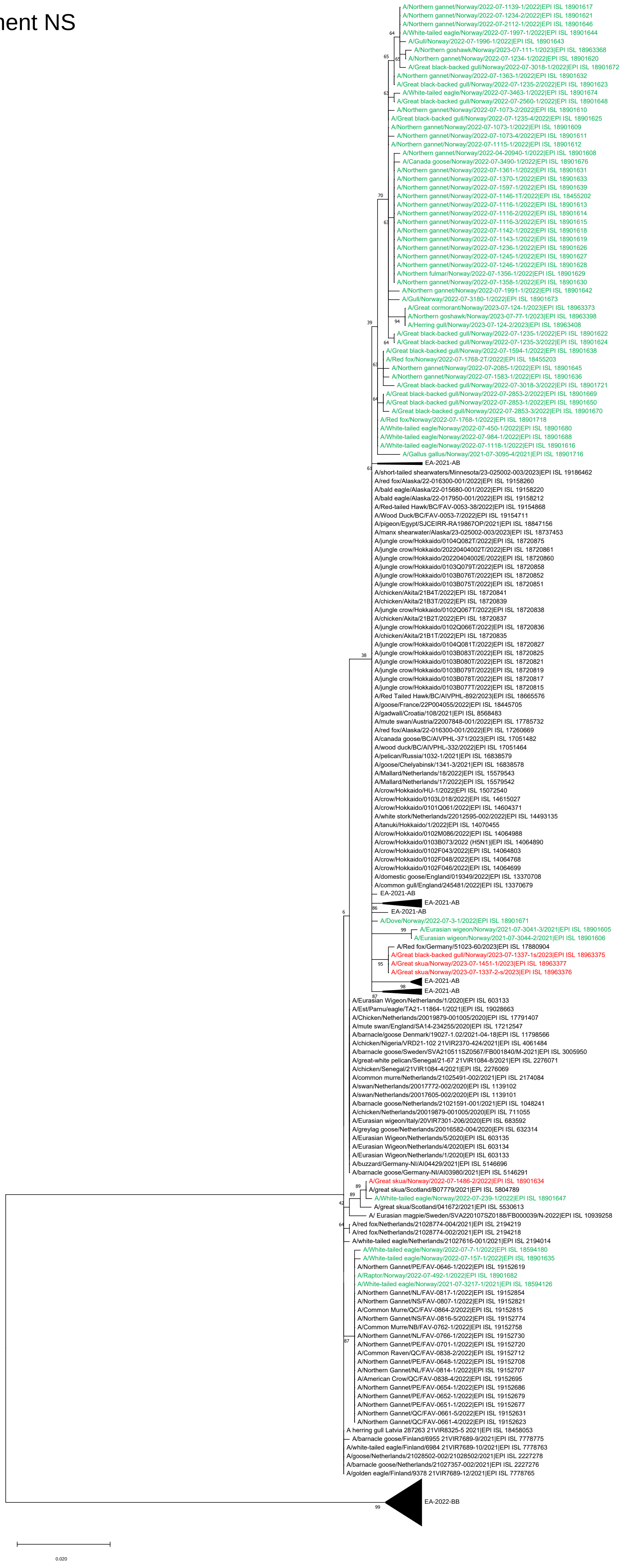

### EA-2021-AB, segment PB2



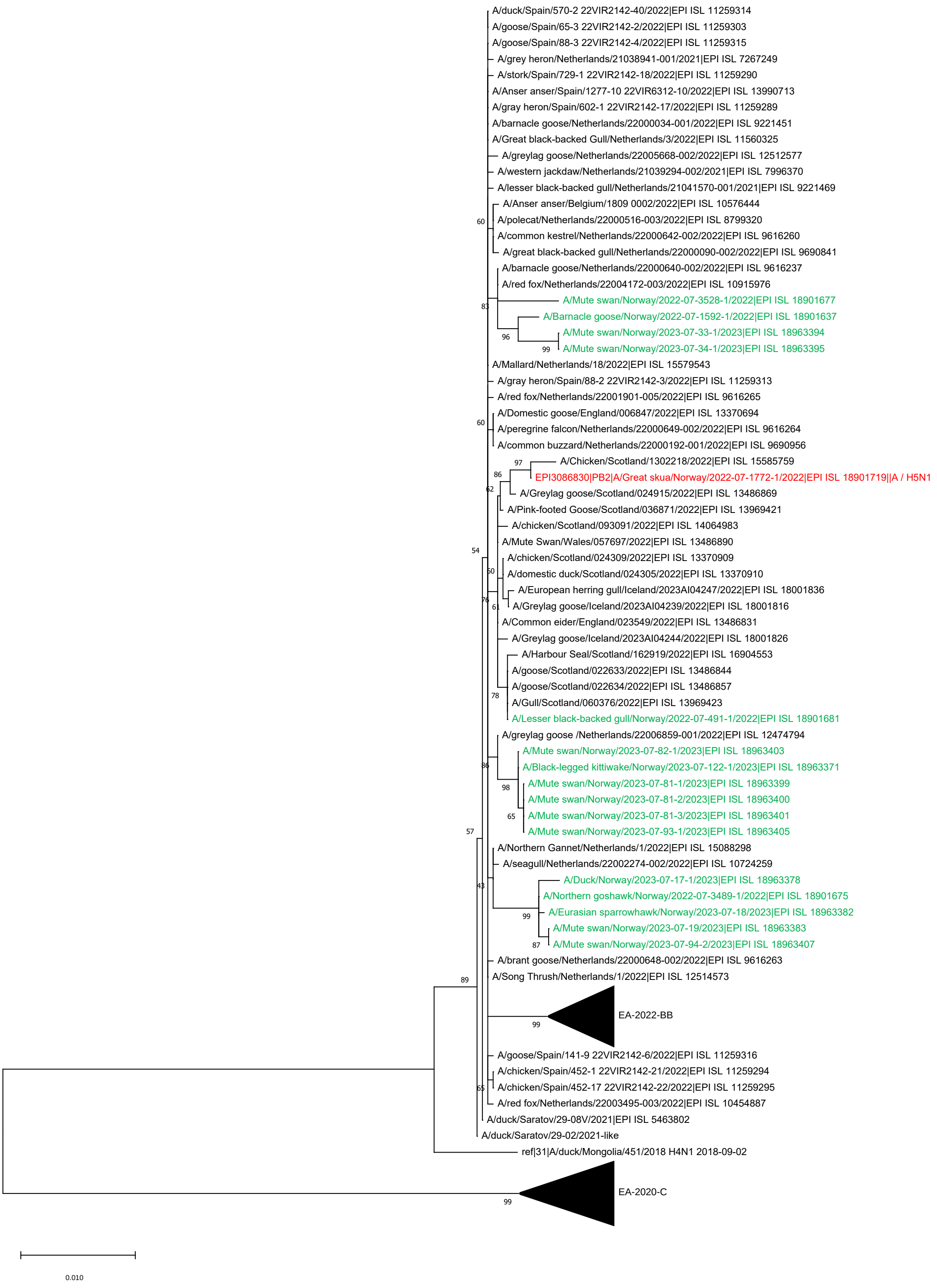

### EA-2021-AB, segment PB1

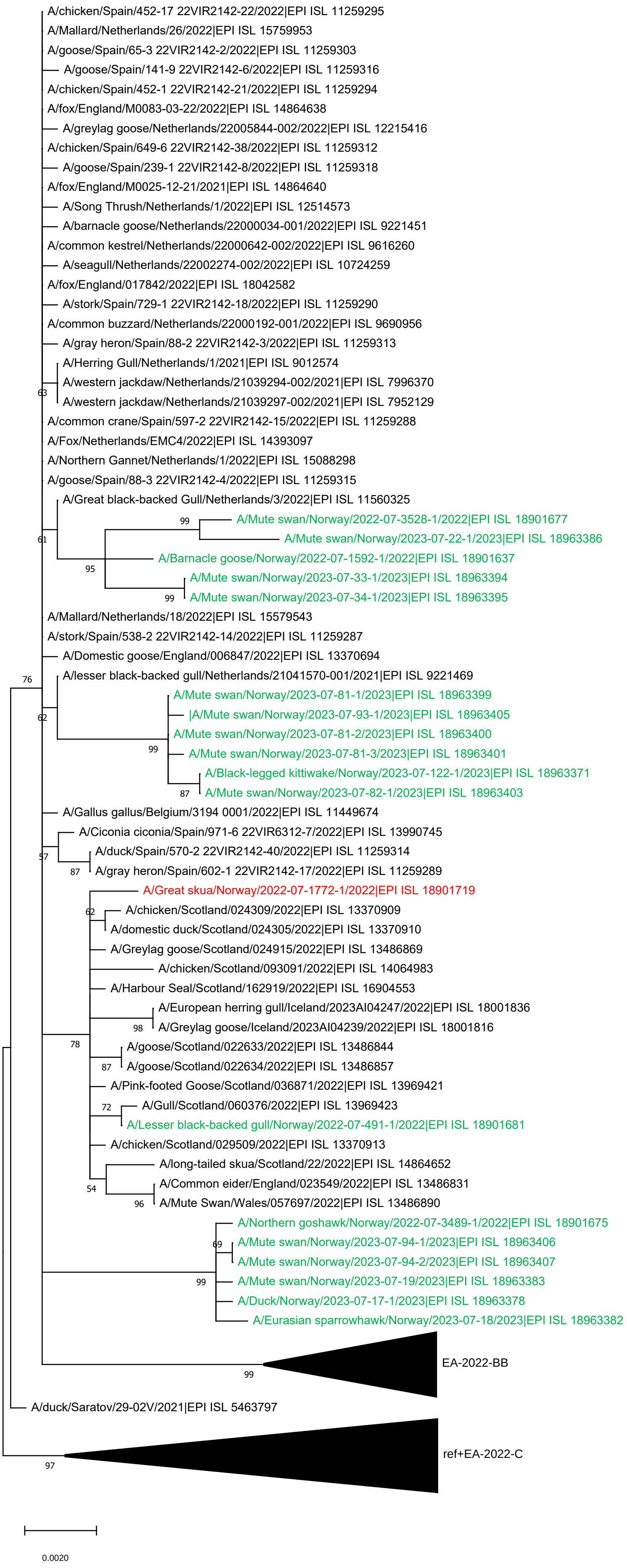

### EA-2021-AB, segment PA

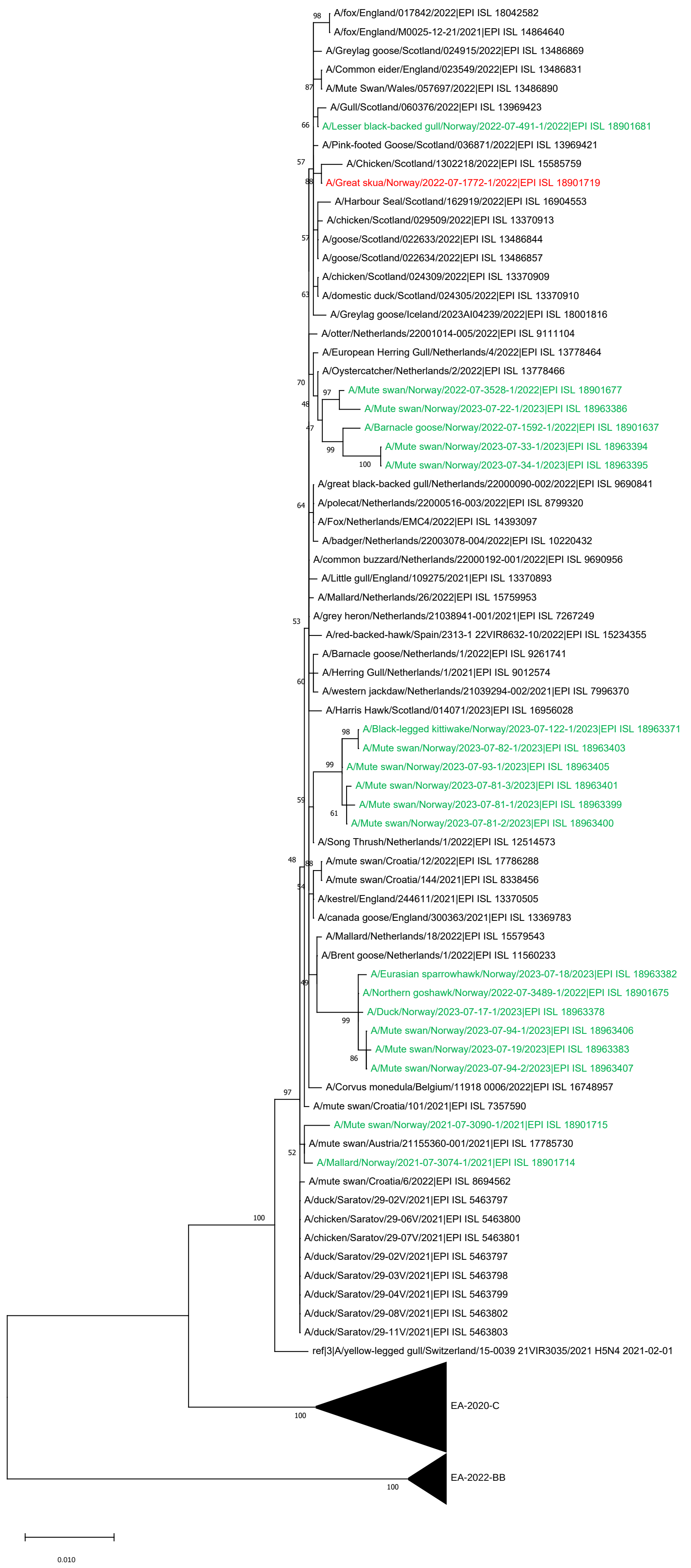

### EA-2021-AB, segment NP

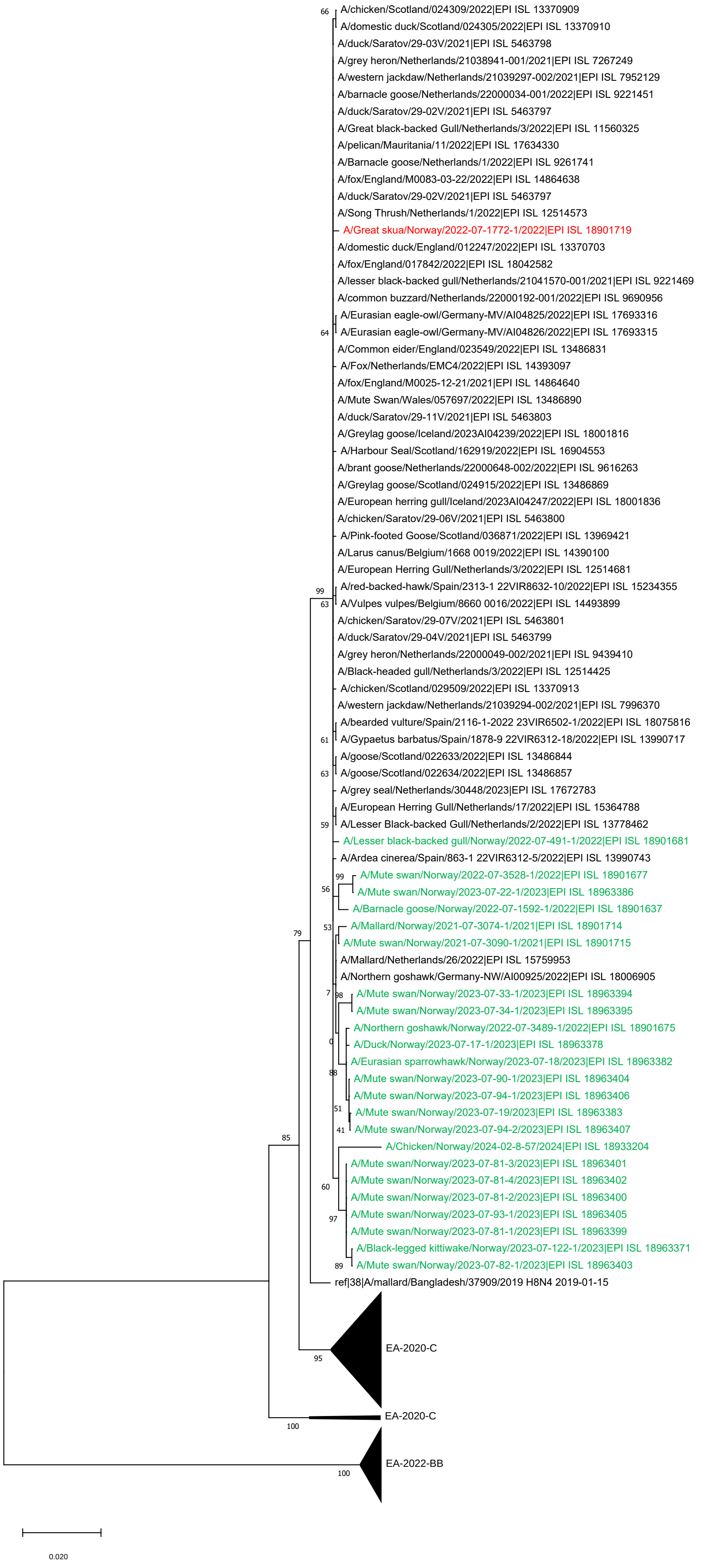

### EA-2021-AB, segment NA

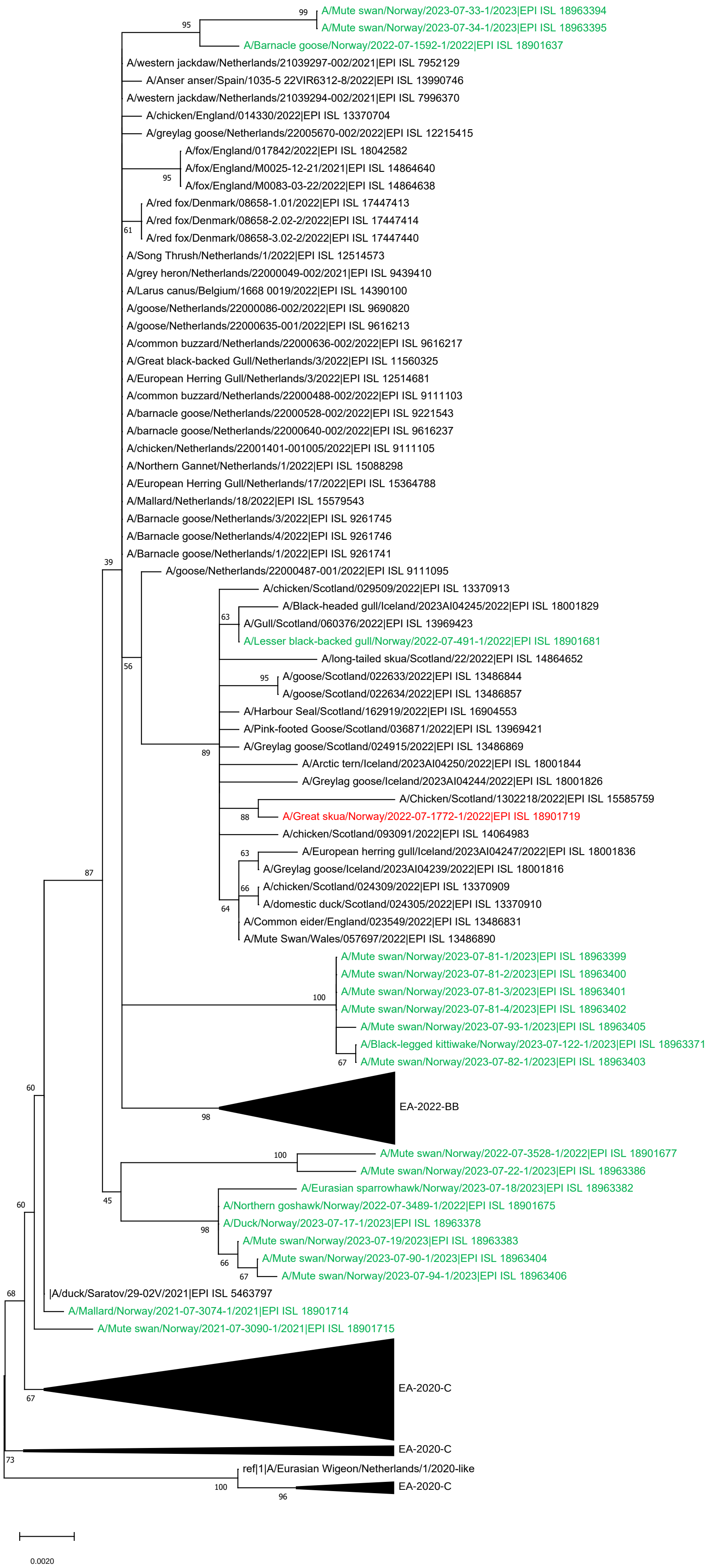

### EA-2021-AB, segment MP

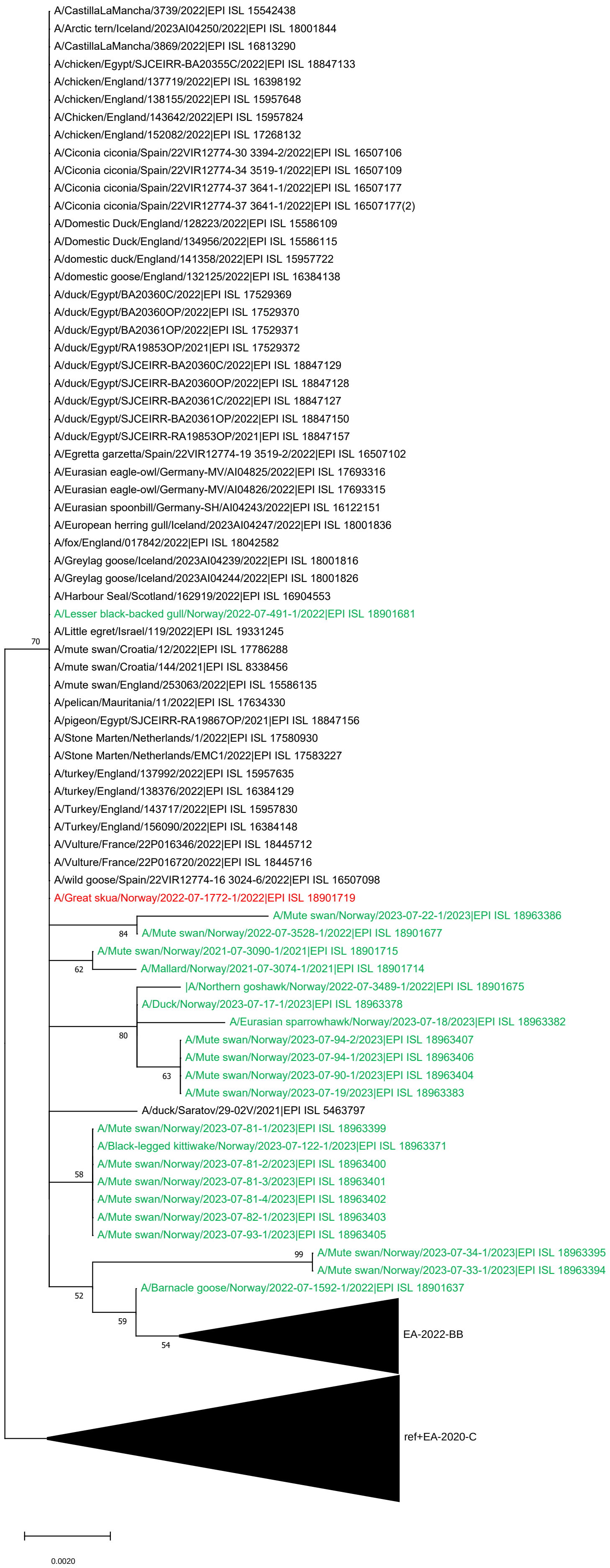

### EA-2021-AB, segment NS

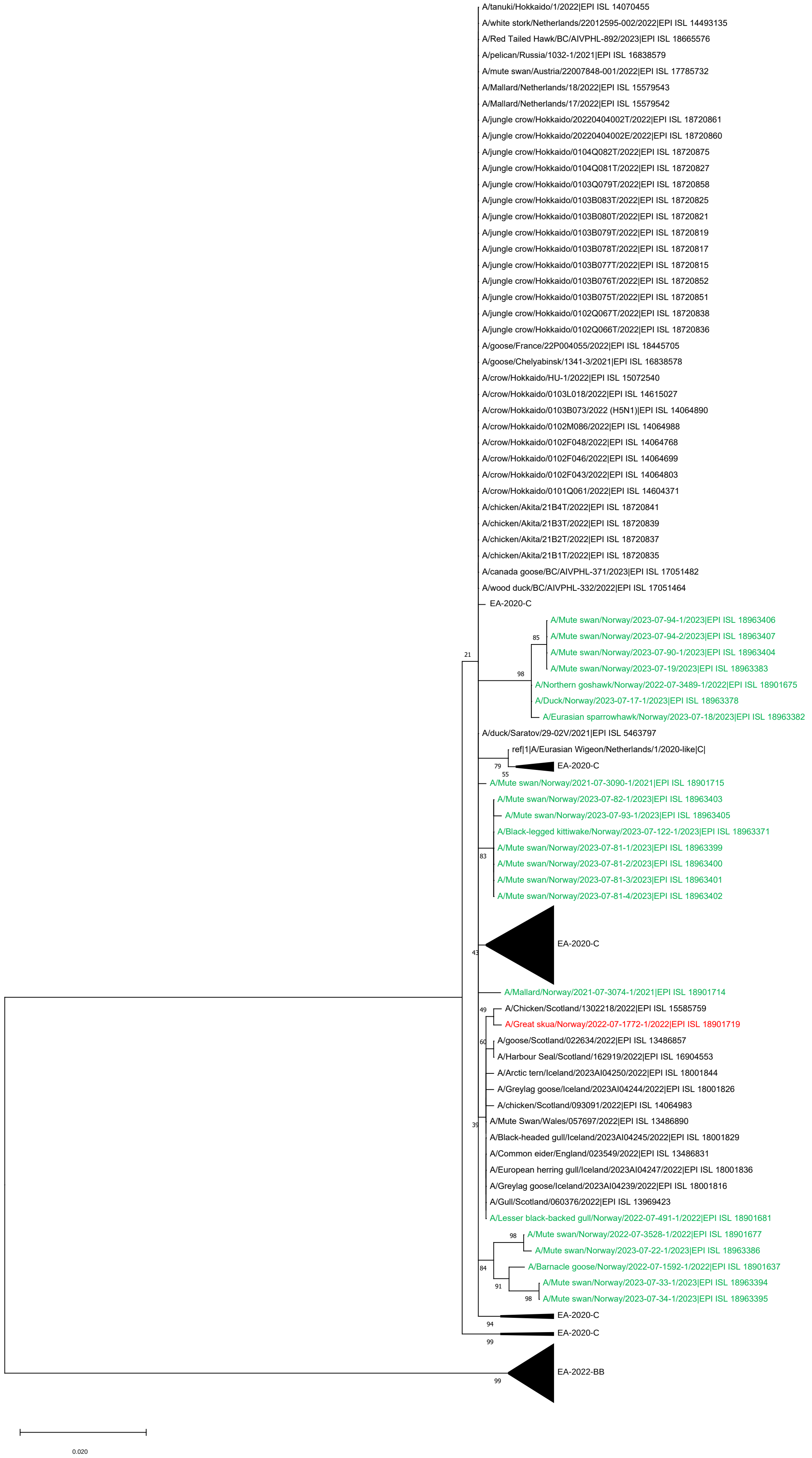

EA-2022-BB, segment PB2

Supplementary Figure 1D

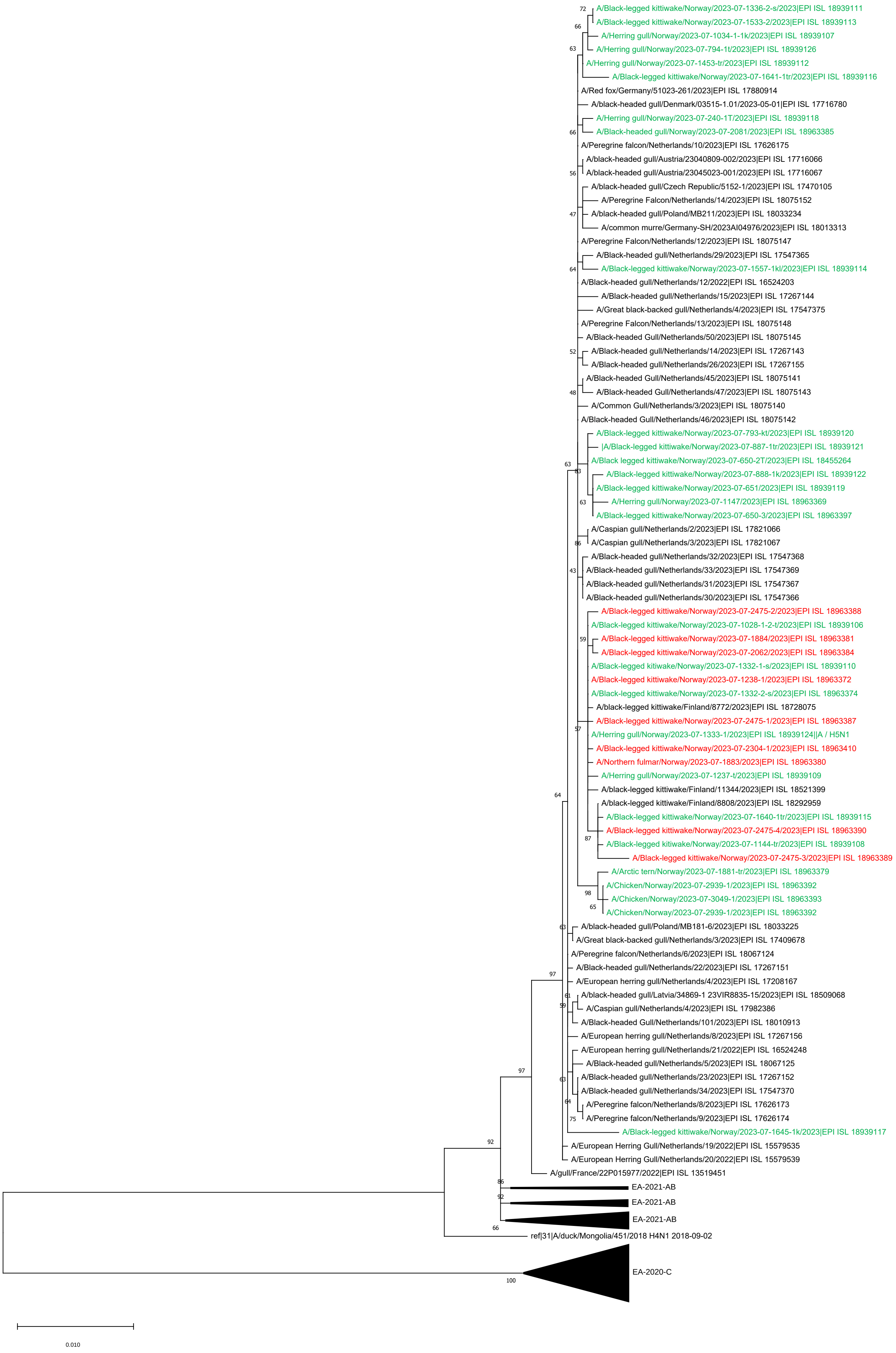

### EA-2022-BB, segment PB1

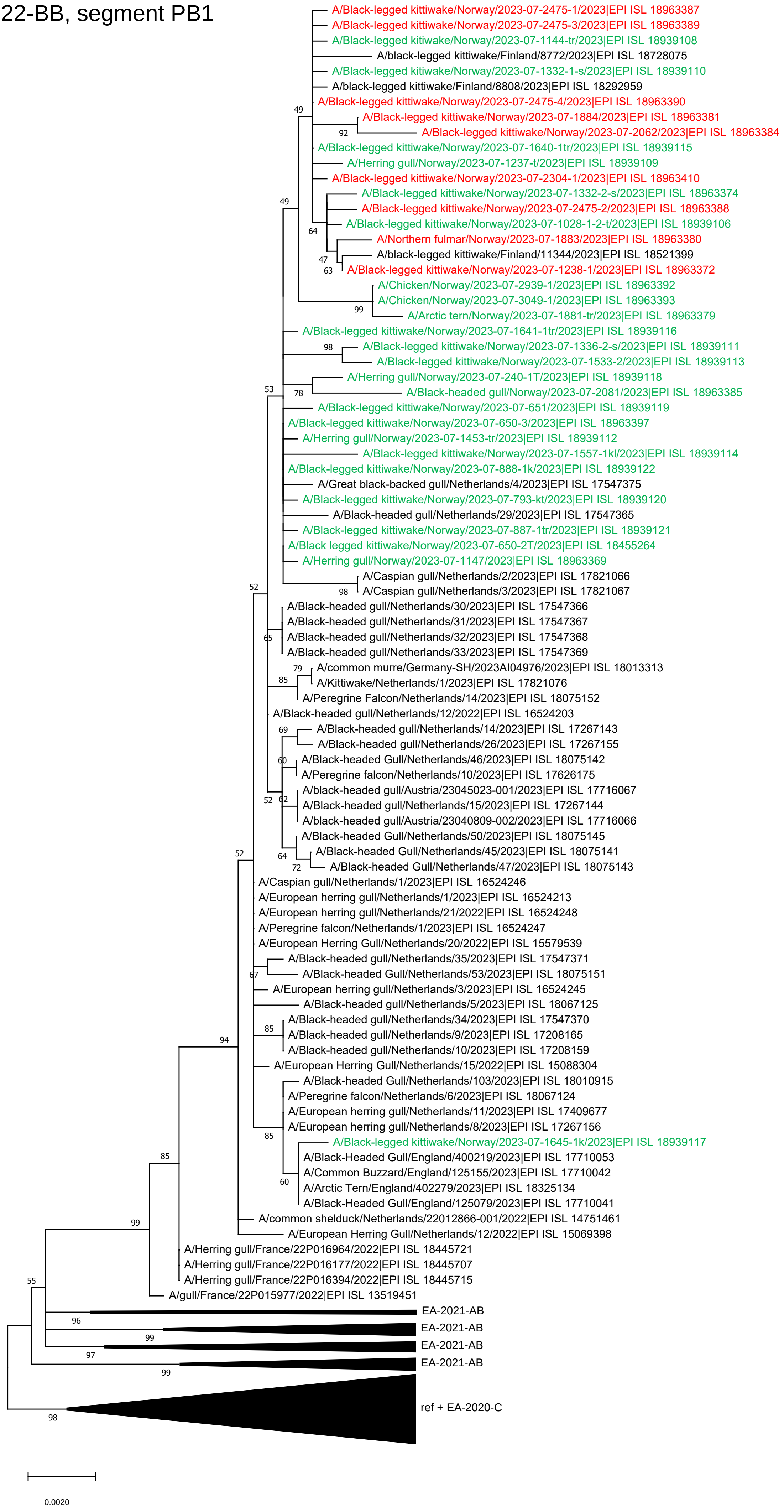

### EA-2022-BB, segment PA

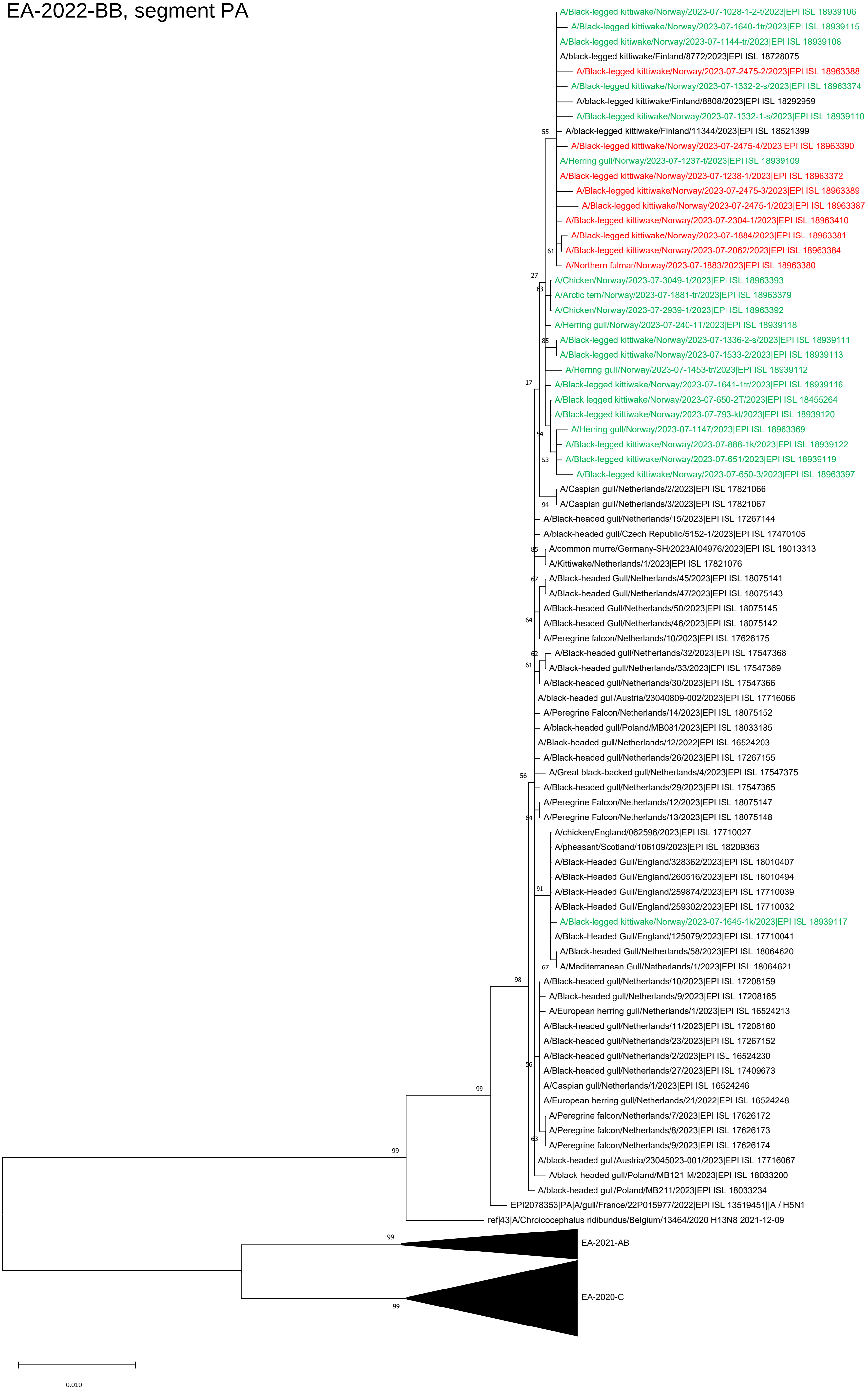

### EA-2022-BB, segment NP

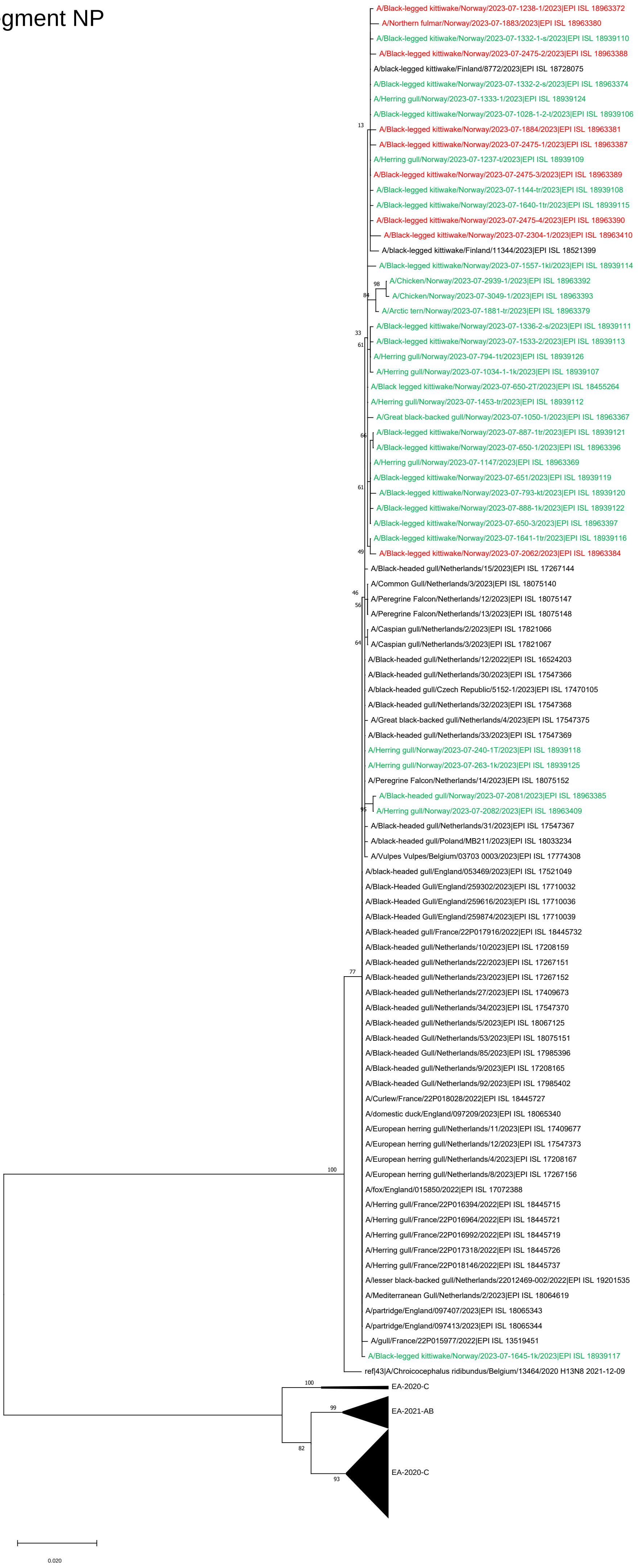

EA-2022-BB, segment NA

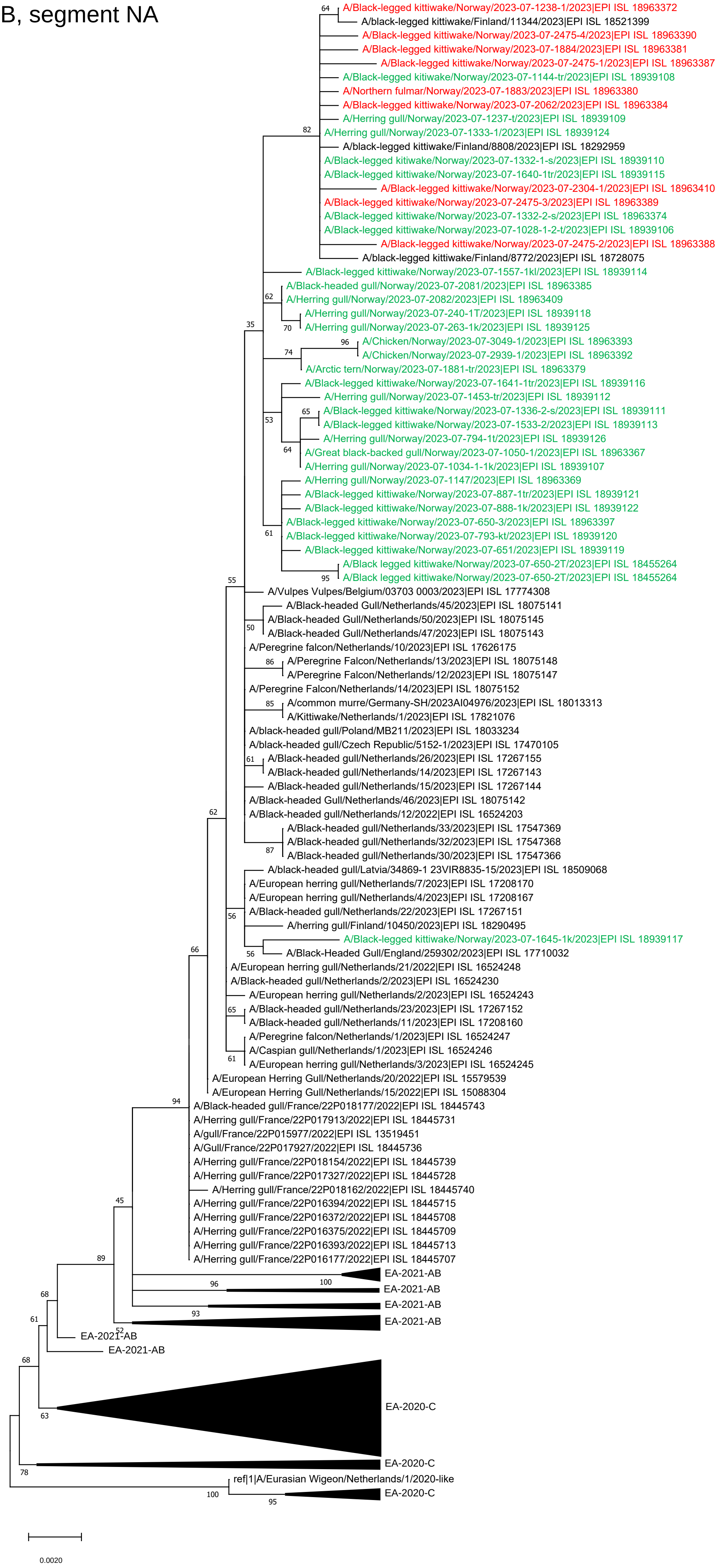

### EA-2022-BB, segment MP

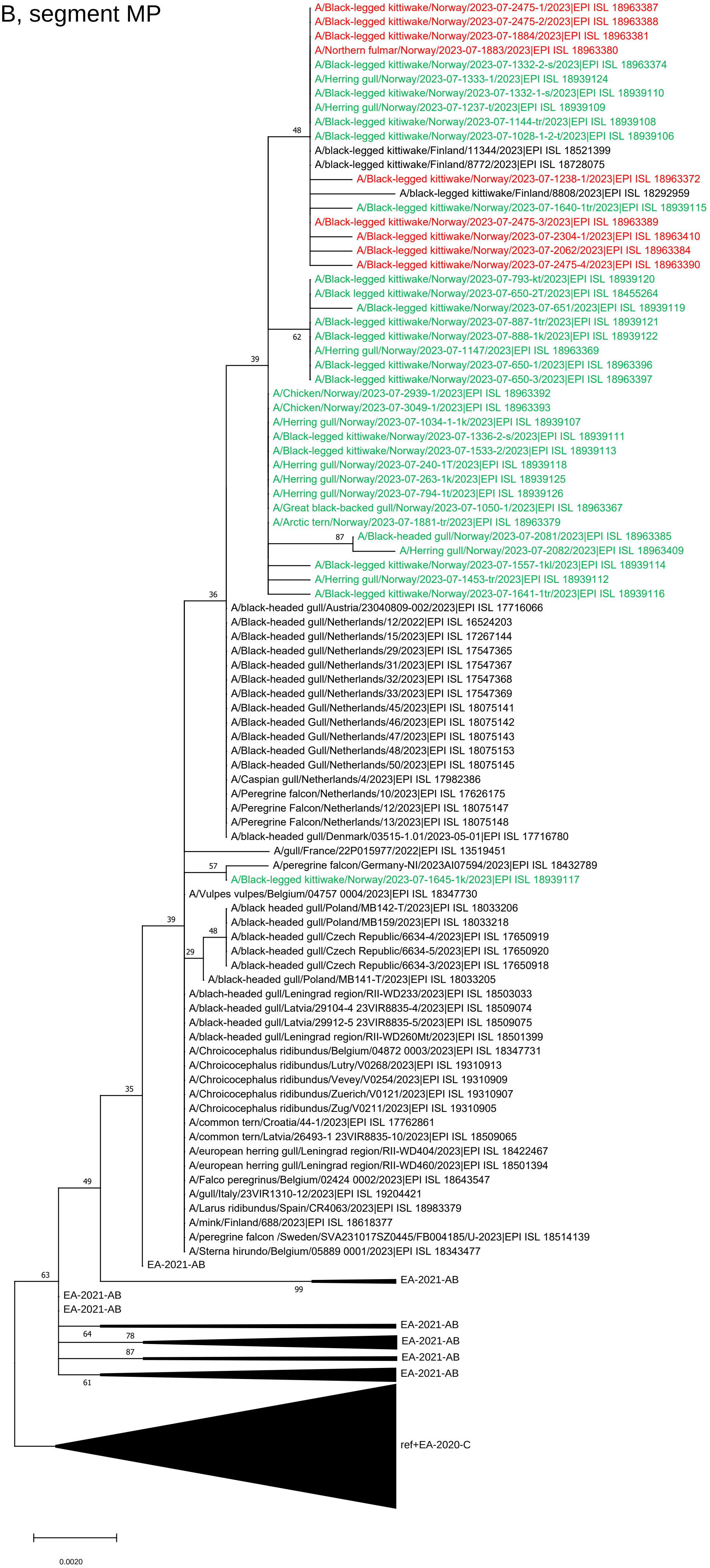

### EA-2022-BB, segment NS

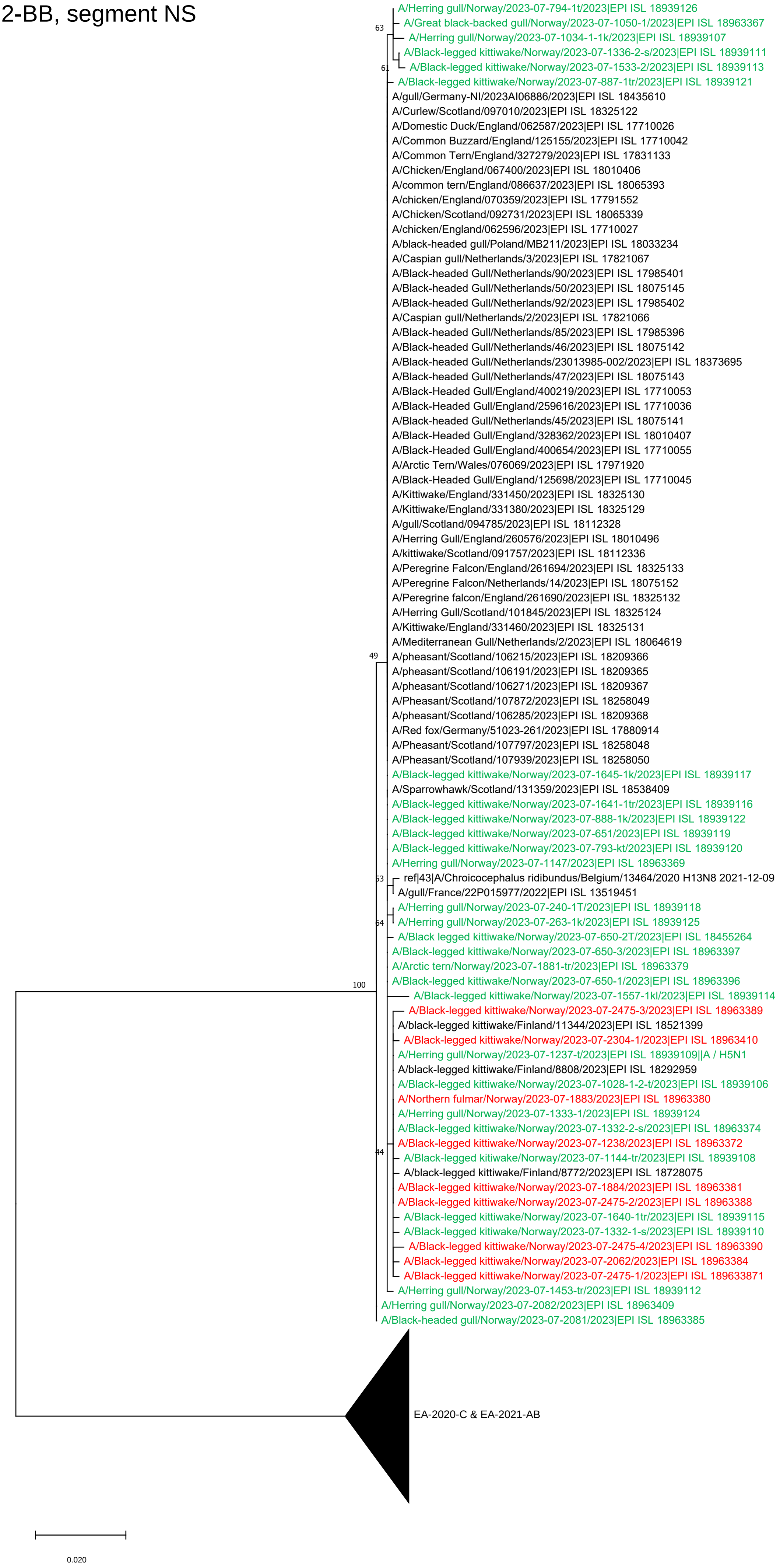
